# Generative models for antimicrobial peptide design: auto-encoders and beyond

**DOI:** 10.1101/2025.10.29.685317

**Authors:** Lukas Beierle, Julian Hahnfeld, Alexander Goesmann, Reihaneh Mostolizadeh, Franz Cemič

**Affiliations:** Bioinformatics and Systems Biology, Justus-Liebig-University Giessen, Ludwigsplatz, Giessen, 35390, Hesse, Germany; Department of Computer Science, University of Applied Sciences Giessen, Gutfleischstrasse, Giessen, 35390, Hesse, Germany; Max-Planck-Institute for Terrestrial Microbiology, Marburg, 35043, Hesse, Germany

**Keywords:** antimicrobial peptides, generative models, deep learning, variational auto-encoders, wasserstein auto-encoder, language models

## Abstract

**Background:** Since the number of multi-resistant pathogens is growing rapidly, new strategies to accelerate the development of antimicrobial drugs are urgently needed. A promising candidate class for new antibiotics are antimicrobial peptides, showing lower tendency to induce antibiotic resistance. High-throughput *in silico* strategies for candidate mining, such as generative deep learning algorithms, have become popular over the last few years and offer novel ways for peptide discovery.

**Methods:** This study presents a comparative analysis of contemporary deep learning models’ generative performance for generating novel antimicrobial peptides. The models examined include Variational Auto-Encoders, a Wasserstein Auto-Encoder, a Recurrent Neural Network and a Language Model. The primary focus of this study is the systematic comparison and evaluation of various methods and sampling options to identify the most suitable model and sampling strategy combination for different use cases.

**Results:** The findings demonstrate the models’ capacity to generate peptide sequences exhibiting analogous properties to those of naturally occurring active peptides, which are utilized for model training while featuring an appropriate degree of sequence diversity. Auto-encoder-based models, particularly the Wasserstein auto-encoder, have generated novel and remarkably diverse sequences compared to recurrent neural networks and language models. This model category exhibits a propensity to prioritize the frequencies of individual amino acids during the learning process, in contrast to variational auto-encoders. Furthermore, latent space models have been shown to possess the capacity to utilize diverse methodologies for generating novel peptides. However, it is imperative to note that these sampling strategies are not universally advantageous or disadvantageous; their optimal selection is contingent on the specificities of each individual use case.

**Conclusion:** The present study investigates the strengths and weaknesses of various generative models for antimicrobial peptides and suggests which model and sampling strategy combination should be favoured for specific individual applications.

## 1 Introduction

It is indisputable that there is an urgent need for the development of novel antibiotic agents and alternative treatment strategies to combat infectious diseases. Since the traditional development of antibiotics is associated with high costs in terms of time and money, new strategies have to be developed [1]. Infections with drug-resistant pathogens remain a significant global health concern, impacting millions of lives annually. Accurately determining the case and death numbers associated with these infections presents a complex challenge due to variations in healthcare quality, reporting standards, and accessibility across different regions.

A recent study examined older forecasts of cases and deaths and incorporated historical trends and future predictions in their data. This study’s findings indicate that, from 2024 until 2050, the number of deaths attributable to Antimicrobial resistance (AMR) could exceed 39 million [2, 3]. Among the most concerning contributors to this crisis are the so-called ESKAPE pathogens [4–6], a group of bacteria known to have a high capacity to develop resistances to standard antibiotic therapies. The term “ESKAPE” encompasses *Enterococcus faecium, Staphylococcus aureus, Klebsiella pneumoniae, Acinetobacter baumannii, Pseudomonas aeruginosa* and *Enterobacter species*. Often *Escherichia coli* is also considered a member of this group. Due to their multifaceted resistance mechanisms, these pathogens are prominent causes of nosocomial infections. These include (i) the production of antibiotics degrading enzymes, (ii) the adaption of efflux pumps that expel antibiotic substances from the bacterial cells, and (iii) modifications in membrane composition [7–10].

Antimicrobial peptides (AMPs) are a particularly promising class of new antibiotics. These peptides are typically short, consisting of 10 to 50 amino acids, and are characterized by their ability to disrupt the membrane integrity of microbial cells, leading to cell death [6, 9]. Furthermore, it is well known that bacteria are less prone to develop resistances against AMPs than classical antibiotics. This is mainly due to the fact that AMPs can act on different hydrophobic and/or polyanionic targets [11–13]. AMPs are found in almost all living organisms, from bacteria to humans, indicating their evolutionary importance as a first defense against pathogenic microorganisms [14]. The mechanisms of action of antimicrobial peptides frequently involve the targeting and permeabilizing of microbial membranes [15], yet they are also capable of interacting with intracellular targets, disrupting vital processes within the pathogens [16]. These multifaceted mechanisms of action render AMPs highly effective against a broad spectrum of microorganisms, including bacteria, fungi, viruses, and parasites. Moreover, AMPs have demonstrated efficacy against antibiotic-resistant strains, underscoring their potential as alternative therapeutic agents in an era of increasing antibiotic resistance [9].

Generative deep learning offers significant advantages in developing novel AMPs as antibiotics. It accelerates the discovery process by rapidly generating diverse peptide sequences, which increases the likelihood of finding effective candidates [1]. Generative deep learning helps designing peptides less susceptible to resistance mechanisms by creating novel sequences. Overall, this cost-effective and efficient approach offers a promising avenue for developing new antibiotics against drug-resistant pathogens. In recent years, there has been great progress in computational biology, with a particular focus on the development of deep generative models. Early efforts in this area relied on Recurrent neural networks (RNNs) to capture sequential patterns and generate peptide-like sequences. Müller *et al*. [17] demonstrated the ability of RNN-based models to generate functional AMPs with desired biological properties.

As generative models evolved, more complex architectures such as variational auto-encoders (VAEs) and Generative adversarial networks (GANs) were introduced [18, 19]. While GANs are capable of generating highly realistic sequences, they are often more challenging to train due to issues like mode collapse and instability in the training process [18, 20]. In contrast, VAEs offer a more stable training framework. Advanced variants of VAEs, including Wasserstein auto-encoders (WAEs), have been shown to generate peptide sequences of higher quality, as demonstrated in the research conducted by Das *et al*. [1]. The majority of studies in this domain currently focus on VAEs [21–26] or advanced variants, such as WAEs [1, 27]. In addition, the model proposed by Szymczak *et al*. [28] introduces an innovative hybrid approach incorporating multi-objective optimization techniques. This technique allows more precise control of peptide properties and activity during training.

The advent of transformer-based models, as described in the seminal research paper “Attention is All You Need” by Google DeepMind [29], has given rise to novel approaches in the domain of sequence generation. These architectures leverage self-attention mechanisms to capture long-range dependencies in sequences, enabling more sophisticated peptide designs with enhanced functional properties.

In addition, there are greedy methodologies, as demonstrated by the work of Wan *et al*. [30], where the authors mined the entire space of peptides of specified length in order to identify new antimicrobial peptides. Furthermore, the increasing availability of diverse datasets from genomics and proteomics has resulted in the development of novel methodologies for identifying AMPs. To illustrate this point, Santos-Junior *et al*. [31] have developed a workflow that facilitates screening genomes and meta-genomes with a specific focus on AMPs. King *et al*. [32] searched the human microbiome for peptide-based antimicrobials. Additionally, Maasch *et al*. [33] have successfully mined the proteomes of extinct animals to generate new antimicrobial peptides.

Here in this study we develop and implement various deep-learning-based methods for antimicrobial peptide generation. In addition, we propose appropriate use cases for the methods developed.

## 2 Methods

### 2.1 Data collection and preprocessing

#### Training data

Active AMP sequences were collected from the following publicly available databases: CAMPR4 [34], DBAASP [35], dbamp [36], PlantPepDB [37], APD3 [38] and LAMP [39]. It should be noted, however, that the sequences from the LAMP database were not collected directly from the database but from the work of Bournez *et al*. [40] because the LAMP web service was no longer available. The date of the last state of the databases is given in supplementary table 2. The collected sequences were filtered according to the following criteria: They were required to contain only canonical amino acids. Their sequence length was required to be between 3 and 36, enabling uncomplicated solid-state peptide synthesis. Furthermore, their minimum inhibitory concentration (MIC) or inhibitory concentration (IC) was required to be ≤ 35 µM, with activity against at least one of the organisms of the ESKAPE group, including *E. coli*. In the case of the auto-encoder-based models, the training dataset was randomly split by 90:10, with the second part being utilized for the reconstruction of known AMP sequences. This distinct dataset can be regarded as the standard test dataset; henceforth, it will be designated as the reconstruction dataset. The RNN and the language model were trained on the entire dataset. The rationale for this decision is that RNNs and Language models (LMs) cannot be used for template-based sequence reconstruction, like auto-encoders.

#### Comparison data

Three datasets were created for comparison of the generated sequences. The first contains sequences from the UniProt [41] database with the following search parameters: The keywords (all linked with not conditions) 0929, 0295, 0044, 0043, 0930, no pharmaceutical or biotechnological use. The protein existence level is not predicted/uncertain, and the sequence length is between 3 and 36. The final dataset encompasses 6.385 sequences, intending to ensure that these sequences do not comprise active AMPs. The second dataset comprises random sequences of equal length and amino acid distribution as the sequences in the training dataset. Care was taken to ensure that the second dataset and the training dataset were of equal size. The third dataset consists of 40.000 totally random sequences with lengths between 3 and 36 amino acids. It should be noted that the final two datasets have been utilized exclusively to compare the outcomes of the prediction of antimicrobial peptides, as outlined in section 3.5.

### 2.2 Sequence encoding

The OneHot encoding scheme was used in all auto-encoder-based models. Prior to this, all sequences were adjusted to a uniform length of 36. Hyphen characters of appropriate length are used as prefixes to extend smaller sequences. According to the model’s documentation on KerasHub [42], a specific padding character was employed in the context of language models. This character was denoted by the token `“[PAD]”`. In addition to this, the token `“[BOS]”` was inserted at the beginning of each sequence to demarcate its starting character. For this purpose, the `StartEndPacker` from the KerasHub library was used. In contrast to the other models, the language model utilized a distinct encoding method. The `TokenAndPositionEmbedding` layer from KerasHub was used for a positional embedding. The training sequences utilized by the RNN were processed following the methodology outlined by Müller *et al*. [17].

### 2.3 Generative models

#### Variational auto-encoders

VAEs, first described by Kingma *et al*. [43], are an extension of classical auto-encoders based on variational inference. These models consist of an encoder *q*_*ϕ*_ and a decoder *p*_*θ*_, each represented by a neural network with parameter sets denoted by *ϕ, θ*, respectively. In a variation inference setting, the objective is to approximate the given dataset’s unknown distribution *p*(*x*). The VAE achieves this by approximating *p*(*x*) with the marginal likelihood *p*_*θ*_(*x*). For this purpose, the encoder maps every datapoint *x* into a continuous latent space *p*(*z*) forced to be standard normally distributed. During training, the VAE aims to maximize the evidence lower bound (ELBO) of the marginal likelihood *p*_*θ*_(*x*), which is equivalent to minimizing the Kullback-Leibler divergence (KL) between the posterior approximated by the decoder *q*_*ϕ*_(*x* | *z*) and the true posterior *p*_*θ*_(*z* | *x*).

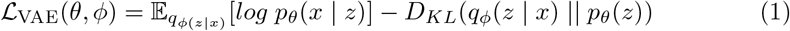

#### Wasserstein auto-encoder

WAEs are an extension of VAEs proposed by Tolstikhin *et al*. [20]. Instead of using variational inference as an objective, WAEs rely on optimal transport *W*_*c*_(*P*_*X*_, *P*_*G*_) where *P*_*X*_ is the actual data distribution and *P*_*G*_ is the approximation by the model. *G*(*Z*) denotes a sample from the decoder. The distance between the input and the reconstructed data is measured by a cost function *c*(*P*_*X*_, *P*_*G*_) chosen as the mean squared error in this work. Instead of computing a pointwise approximation to the latent prior *P*_*Z*_ as is usually done by VAEs, the WAE forces a continuous mixture *Q*_*Z*_ := ⎰*Q*(*Z* | *X*)*dP*_*X*_ to match *P*_*Z*_ [20]. Rather than using KL, the WAE is regularized with a maximum mean discrepancy (MMD) term *D*_*Z*_(*Q*_*Z*_, *P*_*Z*_) in eq. 2 [20, 44]. Specifically, as proposed by Tolstikhin *et al*., [20] we used an inverse quadratic mean kernel for calculating the MMD and introduce *λ* as a scaling-parameter. This leads to the optimization problem given by eq. 2.

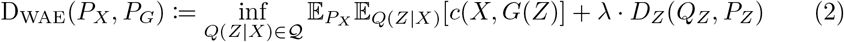

#### Recurrent neural network

As described by Müller *et al*. [17], RNNs were initially used in natural language processing and can capture medium-range dependencies in sequences. Later on, more advanced variants such as Long-Short-Term-Memory (LSTM) cells [45] and Gated-Recurrent-Units (GRUs) [46] were developed to overcome certain limits of RNN such as vanishing or exploding gradients [17, 45]. GRU units were developed to simplify LSTM cells. RNNs can also be used to generate new sequences. For this, one or more starting characters can be provided to the model. Then, it recursively predicts the most probable characters based on the representation learned during training. As an objective function, the RNN uses categorical cross-entropy.

#### Language model

LMs are a type of architecture based on the seminal paper entitled “Attention is all you need” by Google DeepMind [29], that introduces the transformer architecture into natural language modeling. Various implementations designed for protein sequence generation were published [47–49]. Inspired by the architecture of ProtGPT-2 [49], a transformer-based decoder-only model, a small language model for AMP sequence generation was also developed in this study. Language models typically undergo extensive training on very large datasets, often with prolonged pre-training periods requiring substantial resources. In contrast, the present study uses a smaller dataset, leading to the decision to implement a compact language model. This model has a single `TransformerDecoder` layer with two attention heads, an embedding dimension of 32, an intermediate dimension of 32, and a dropout rate of 0.1. The LM’s sampling procedures differ from those of the RNN, VAEs, and WAE. The LM uses its own sampling algorithms, provided by the KerasHub library. LMs generate new sequences based on the probabilities of amino acids during training. The samplers select the next amino acid for the generated sequence.

### 2.4 Model architectures and implementations

The models introduced in section 2.3, namely a WAE, an RNN, and four distinct VAEs featuring different regularization schemes, were implemented together with a small LM. Figure 1 shows a visualization of the different model architectures. The RNN comprises two interconnected LSTM layers, with 256 units each, followed by a dense layer with 21 units and the softmax activation function. The encoder’s architecture comprises an input layer, two GRU layers with 128 units, a batch normalization layer, an attention layer that performs self-attention, three 1D-convolution layers, and two dense layers representing *µ* and *σ*. Notably, in the case of the WAE, there is a single dense layer instead of two. The decoder is composed of an input layer, followed by a `RepeatVector` layer, which is employed to ensure that the appropriate input dimension is provided to the subsequent two GRU layers with 128 units each. A dense layer comprising 21 units, combined with the softmax activation function, is utilized as the final layer. A summary of the used hyperparameters: batch size, number of epochs, learning rate and the size of the latent dimension can be found in supplementary table 3. It is important to note that the learning rate of the WAE was reduced with an `ExponentialDecay` schedule from the Keras library [50] during training, which was also done in the original work of Tolstikhin *et al*. [20].

**Fig. 1.**
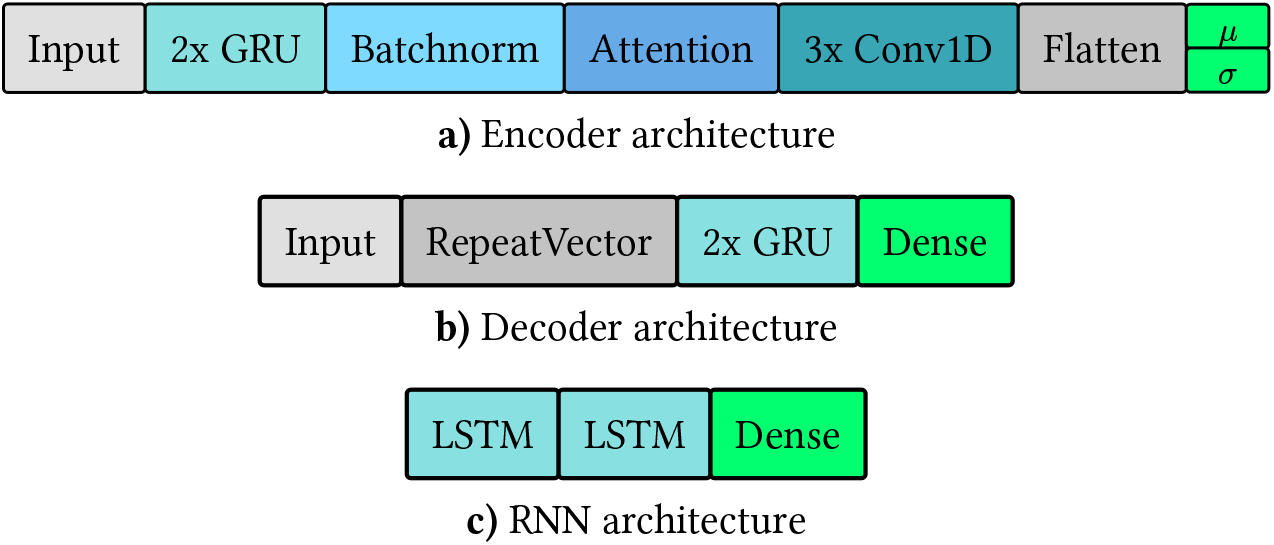
Visual representation of the models’ architectures. Architectures used for the encoder and decoder of the VAEs, WAE, and the RNN. **a)** The encoder has an input layer, two GRU layers and a batch normalization layer. Next, an attention layer performs self-attention. Then, three 1D convolution layers are used, ending in a flatten layer with an additional dense layer. In the case of the WAE there is only one dense layer. **b)** The decoder’s architecture comprises an input layer, a RepeatVector layer to match the required input shape of the two consecutive GRU layers and a dense layer. **c)** The RNN consists of two interconnected LSTM layers and a dense layer.

#### Annealing schedules

A well-known problem encountered during the training of VAEs is the so-called posterior collapse [51]. This problem occurs when both terms on the right-hand side of the equation 1 are of different magnitudes. In the scenario that the reconstruction term is assigned a greater weight compared to the regularization term, the decoder tends to neglect the latent variables and reconstructs the input data directly. Consequently, the KL divergence approaches zero, thereby enabling the maximization of the ELBO. However, this approach prevents the models from learning a meaningful data representation. As a result, the generated data exhibits high similarity to the input data and lacks diversity.

To avoid the VAE-models’ tendency to posterior collapse, KL annealing was used. Specifically, a scaling factor *β* is introduced to the KL term of the objective function in equation 1. The parameter *β* was iteratively modified during training as detailed by equations 3, 4 and 5. In addition, a fixed *β* = 1 (eq. 6) was included for comparison to a non-regularized VAE. The KL-regularization during training forces a VAE to assign greater weight to the reconstruction term from equation 1.

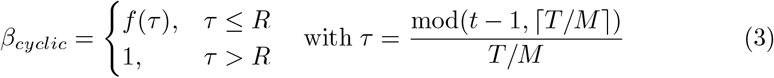

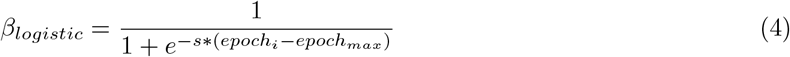

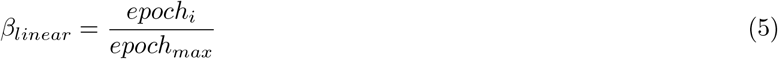

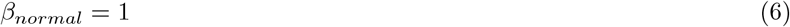

In eq. 3, *t* denotes the current training iteration number, a.k.a. epoch, *T* is the number of epochs, *f* is a monotonously increasing function, while *M* is the number fo annealing cycles during training and *R* is the proportion that is used to increase *β* within one cycle. *M* and *R* were set to the default values as in [52] (*M* = 4, *R* = 0.5). In eq. 4, the parameter *s* controls the steepness of the logistic curve, while *epoch*_*max*_ sets its infection point and *epoch*_*i*_ denotes the current training iteration. The parameter *s* was set to 0.01, while *epoch*_*max*_ was equal to 350.

#### Baseline model

To the best of our knowledge, the RNN from the work of Müller *et al*. [17] was the first model of this type used to generate antimicrobial peptides. The model and training data were made entirely available to the public, enabling full reproducibility. Therefore, the RNN has been selected as the baseline model for this investigation. Later, the VAE models are referenced by their annealing schedule, as shown in the following example: Cyclic (VAE-CYC), linear (VAE-LIN), logistic (VAE-LOG) and without annealing (VAE-N).

The models were trained on a Dell OptiPlex 7090 computer with an Intel Core i7-11700, 32 gigabytes of DDR4 RAM, and an NVIDIA GeForce GTX 1660 Super graphics card. As an operating system, the Linux distribution OpenSUSE Tumbleweed was used.

### 2.5 Sampling strategies

Different methods were used to generate putative new AMP sequences depending on the respective model.

#### Random sampling

All models are capable of generating sequences randomly. In auto-encoder-based models, random sampling refers to drawing a random sample of dimension *sequences* × *length* from the latent prior. The output is then mapped to the space of peptide representations by the decoder and translated to a sequence of amino acids. Random sampling by the RNN is initiated by feeding in a random start amino acid, to which further ones are added recursively until a padding character is emitted or the maximum sequence length of 36 is reached. The cardinality of the set of sequences generated by random sampling was set to 40,000 for all models except the RNN, which was constrained to 7,000 sequences due to its comparatively slower processing speed.

#### Template-based sequence reconstruction

The ability of template-based sequence reconstruction is a feature unique to auto-encoder-based models used in this work. Sequences from the reconstruction dataset served as templates for the reconstruction. Following the training of the VAEs and the WAEs, these sequences are first OneHot encoded and then mapped into the latent space by the encoder. These representations are then fed into the decoder and translated to amino acid sequences. Using that scheme, sequences similar to the templates in terms of amino acid composition and peptide properties are generated, as will be discussed in section 3.

#### Template-based multi sequence sampling

As mentioned, VAEs approximate the latent prior *p*(*z*) point wise. Consequently, if the local approximation is sufficiently precise, multiple samples of a single sequence can be generated by repeating the reconstruction procedure as visualized in figure 2. This sampling strategy is particularly interesting if the template sequence is considered a lead sequence, such as one with desired properties for which different analogs should be generated. The strategy is contingent upon the assumption that the local approximation of the ELBO of the targeted peptide is sufficiently rich. In the absence of this condition, there will be minimal difference in the resulting samples; alternatively, the samples will be identical. For each sequence in the reconstruction dataset a batch of 25 replicates was mapped into the latent space by the encoder. As a result, all sequences were mapped to a local distribution as indicated schematically by the red circle in figure 2. In the next step, a batch of equal size was draw from the latent representation, leading to the reconstruction of the template sequence.

**Fig. 2.**
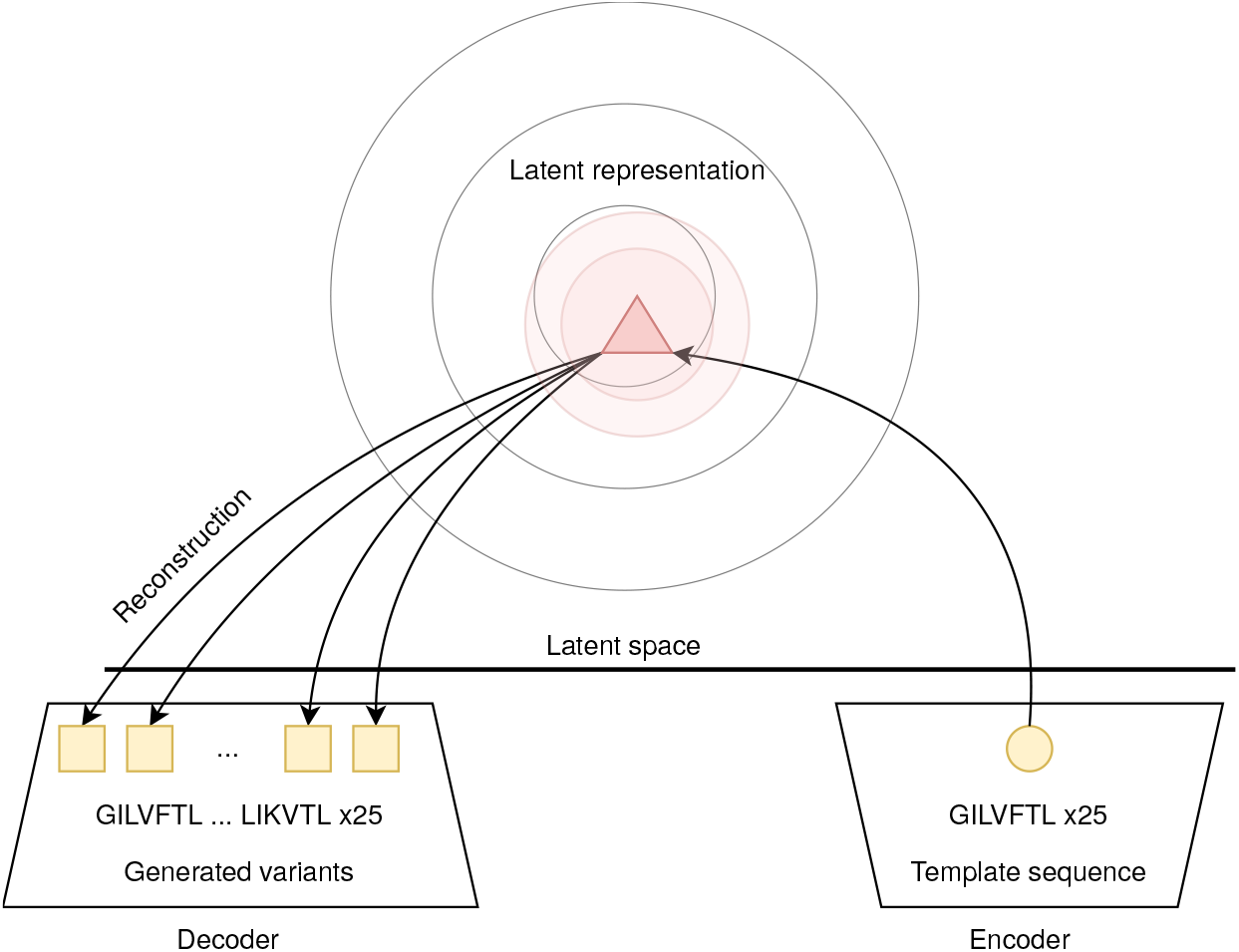
Visual representation of template-based multi sequence sampling. The exemplary input sequence GILVFTL was encoded and repeated 25-times, then mapped into the latent space by the encoder. Since the VAE does a pointwise approximation to the ELBO the 25 sequences are mapped to a distribution highlighted in light red. After this the decoder reconstructs 25 individual samples from the latent distribution resulting in different generated sequences called variants.

**Fig. 3.**
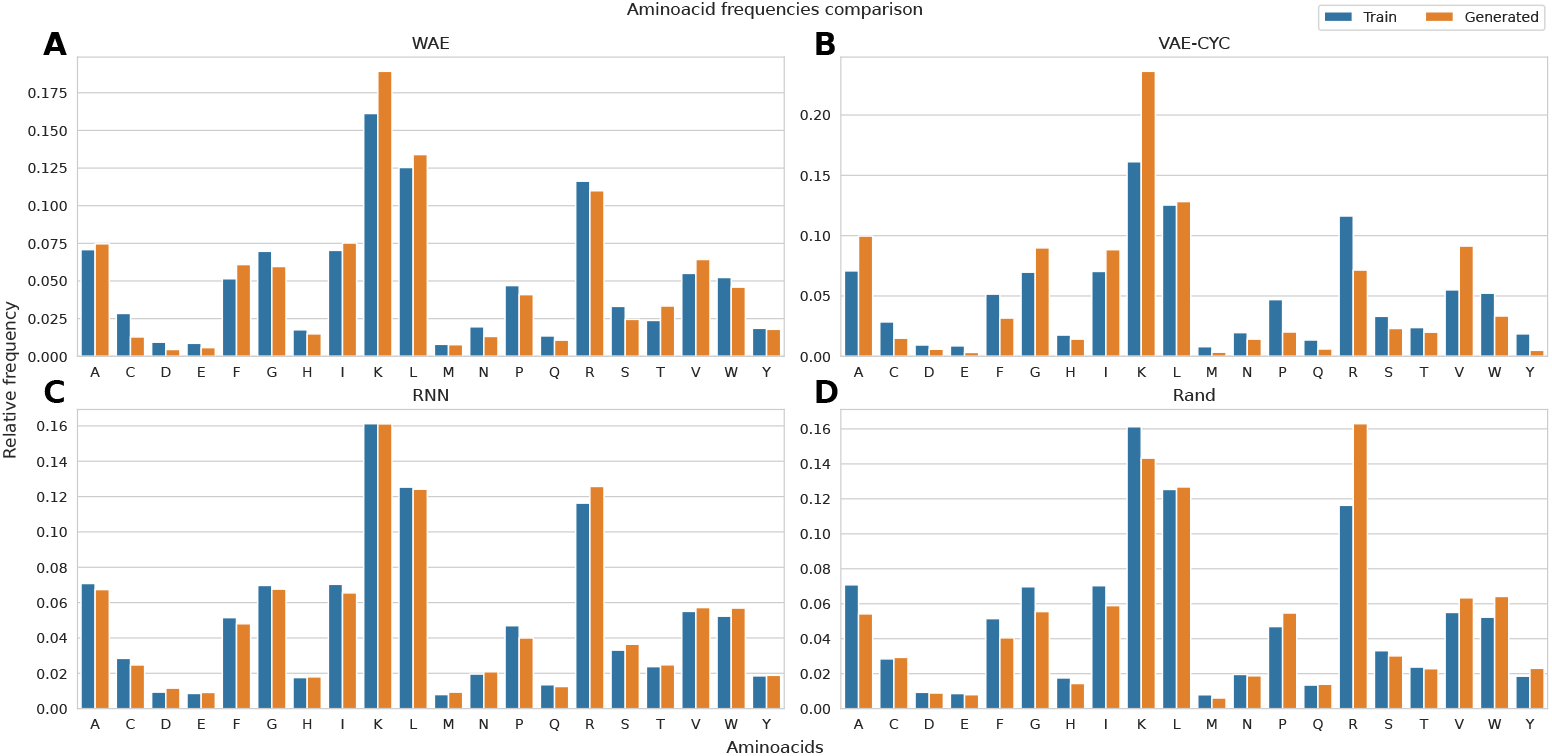
Amino acids comparison between the training set and generated sequences. Each of the sub figures shows the relative frequency of individual amino acids within the training sets (blue) and various generated sequences (orange). The results for the WAE (A), VAE with cyclic annealing (B), RNN (C) and RandomSampler (D) are presented from the upper left to the lower right.

#### Language Model Samplers

The LM utilizes three distinct sampling algorithms in order to generate sequences. These algorithms append the next amino acid with the highest probability calculated by the sampler to the sequence, starting from the start token, and new sequences are generated continuously. This process is then repeated. Three samplers from the KerasHub library were used. The `TopPSampler` with *p* = 0.9, `TopKSampler` with *k* = 10 and a temperature of 1.2. Parameter optimization was performed using gridsearch. For the `RandomSampler`, standard parameters were used.

### 2.6 Sequence selection and evaluation

Generated sequences were filtered according to the following criteria: a sequence length between 3 and 36, the absence of padding characters, and no recurrence of individual amino acids more than seven times. For all valid sequences, training, and comparison datasets, the descriptors of hydrophobicity (Eisenberg scale), hydrophobic moment (Eisenberg scale), charge at pH 7, the isoelectronic point, length, and the mass to charge ratio were calculated using the *peptides*.*py* [53] library in Python. These descriptors were selected based on their correspondence to the activity of AMPs. Furthermore, they have been utilized in previous studies [1, 17]. As illustrated in Figure 4, the distribution of the hydrophobic moment is presented as it is considered the most important descriptor. The figures for the distributions of the remaining data can be found in the supplementary section. At this moment, drawing parallels with other studies concerning the peptide characteristics of generated sequences was not feasible. This was due to an inconsistent data foundation, non-disclosure of models and data, and an absence of reproducibility. In order to facilitate a comparison of the ability of auto-encoder-based models to reconstruct sequences, the reconstruction accuracy per peptide position was calculated according to the method described by Renaud *et al*. [27]. The reconstructed sequences were not filtered, as otherwise, it would no longer be possible to assess the accuracy of the reconstruction.

**Fig. 4.**
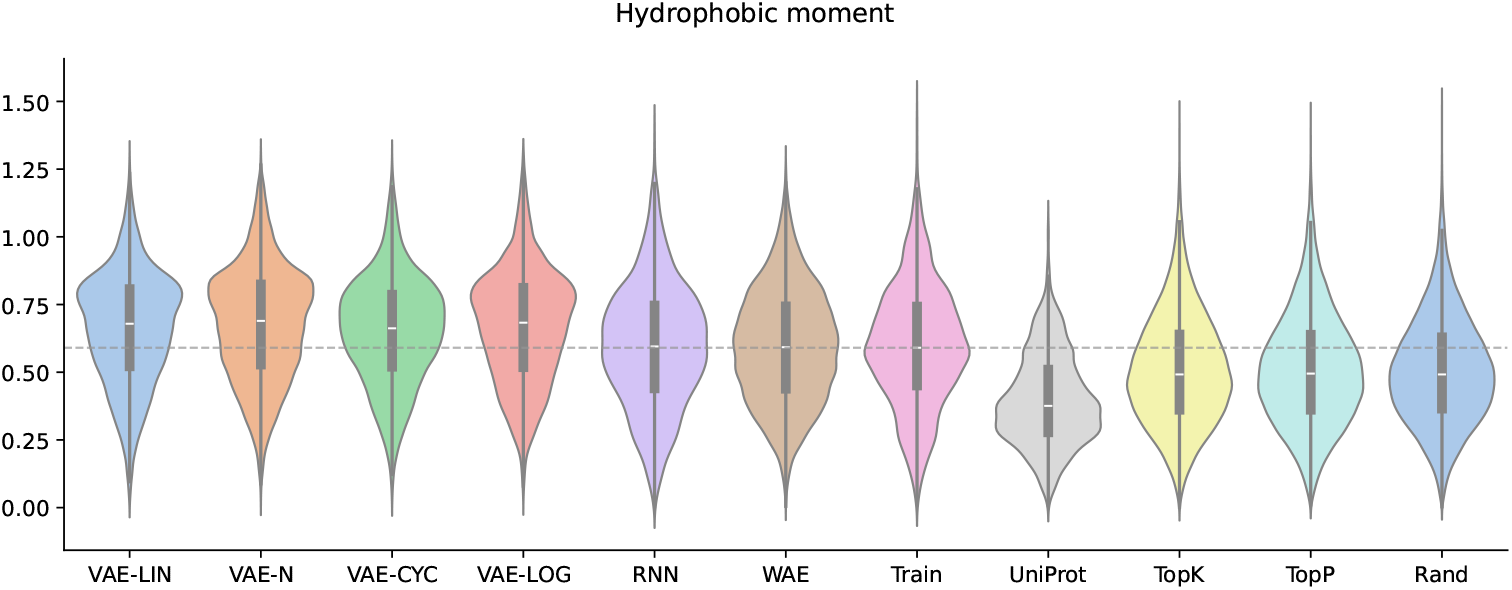
Distributions of the hydrophobic moment. Distributions of the hydrophobic moment shown as violin plots including median and interquartile distances. The distributions of all randomly generated sequences, as well as training and comparison datasets are shown. The gray dashed line marks the median of the training datasets’ distribution for easier visual comparison.

### 2.7 AMP prediction

In order to facilitate further identification of possible active AMPs in the generated datasets, two publicly available tools for AMP prediction were utilized. Firstly, the neural network-based *Antimicrobial Peptide Scanner v2* from the work of Veltri *et al*. [54], and secondly, the recently published tool AntiBP3 from Bajiya *et al*. [55]. Bajiya *et al*. compared their models to other popular publicly available models for AMP prediction and found that their models perform best on independent bench-marking datasets. The model from Veltri *et al*. was also included because the model and datasets are entirely available to the public, and the model can be accessed easily via a web interface. Because both tools necessitate disparate sequence input lengths, all sequences were filtered to a minimum length of 10 amino acids. The results are shown in table 3. The tools mentioned above were chosen for AMP activity prediction because all data relevant for reproducing and testing the model are publicly available.

### 2.8 Manifold learning

In order to compare the peptides generated by the WAE in terms of their molecular properties against known AMPs from the training set, a number of peptide properties, related to AMPs and general ones, were calculated followed by dimensional reduction and visualization. The complete list of calculated peptide properties can be found in supplementary table 5. The molecular weight descriptor was normalized by the length of the peptide to avoid length-related biases. All peptide properties were calculated with the peptides.py library [53]. For dimensional reduction UMAP [56] and t-SNE [57] were utilized. The implementation for UMAP from the umap-learn [58] library was used. For t-SNE we used the implementation provided by the opentsne library [59]. The resulting 2D-embeddings were visualized using the bokeh library [60]. In order to find a trade-off between local and global representation after the dimensional reduction some parameters were adjusted for UMAP and t-SNE. The exact parameters can be found in supplementary table 6.

## 3 Results

### 3.1 Random sequence generation

In this section, the sequences generated randomly are analyzed regarding amino acid composition and peptide properties associated with AMP activity. Figure 3 shows the amino acid composition for the WAE, RNN, VAE-CYC, and the LM with the `RandomSampler`. A summary of mean, maximum, and minimum values and the most prominently deviating amino acid is shown in supplementary table 4. Regarding variations in amino acid composition, the WAE demonstrates the least deviation among the auto-encoder-based models. Compared to the other auto-encoders, this model shows reduced disparities in amino acid composition for less abundant amino acids. Among the VAE-based models, the VAE-CYC features the least deviations, except the mean deviation. In this case, the VAE-LOG model exhibits a smaller value but has higher maximum and minimum deviations. Comparing the `RandomSampler` with the other two samplers reveals that the former exhibits a remarkably lower degree of deviation of amino acid composition. The RNN, on the other hand, shows an almost identical amino acid composition for generated and training sequences. This is due to the modest size of the training dataset, in this case, approximately 7.000 sequences, combined with the fact that no regularization was added for this model. The plots for the other models can be found is supplementary figure 4 and 5.

Figure 4 shows the hydrophobic moment for the generated sequences, the training set, and the UniProt comparison dataset via violin plots. The median of the training set is shown as a gray dashed line for comparison. As demonstrated in figure 4, an upward shift of the mode positions of the hydrophobic moment distributions is evident for the VAE-LOG, VAE-N, and VAE-LIN models. This phenomenon can be attributed, at least in part, to the overrepresentation of the amino acids K and L. It is noteworthy that cyclic annealing serves to mitigate this effect to a certain extent. The WAE and the RNN demonstrate high accuracy in reproducing the median of the peptide property distributions of the training data. However, in the case of the LM samplers, a clear preference for the basic amino acid R is observed, resulting in a lower median of the peptide properties compared to that of the training data. The median of the UniProt reference dataset is below that of the AMPs used for training, which can be attributed to the filters applied in section 2.1 to remove potential AMPs from the UniProt dataset.

Figure 5 shows the charge at pH 7 for the generated sequences, the training set, and the UniProt comparison dataset. The median line of the training set is shown as before in figure 4. The charge distributions of the generated sequences by the VAEs are all shifted slightly upwards compared to the training dataset. This can partly be attributed to an overrepresentation of the positively charged amino acid K. The charge distribution of the sequences generated by the models RNN and WAE closely matches that of the training dataset, with the RNN reproducing the distribution almost perfectly. In the case of the WAE, the median is up-shifted by approximately one electron charge unit. As expected, the distribution of the UniProt dataset is shifted towards lower charge values because most peptides in this set are not expected to interact strongly with bacterial membranes. The distributions of the sequences generated by the different LM samplers are almost identical, with their medians slightly up-shifted compared to the training dataset.

Violin plots for the peptide properties isoelectric point, hydrophobicity and length, together with the amino acid composition diagrams for VAE-L, VAE-N, VAE-LOG and the TopK and TopP LM samplers can be found in supplementary figures 1-5.

**Fig. 5.**
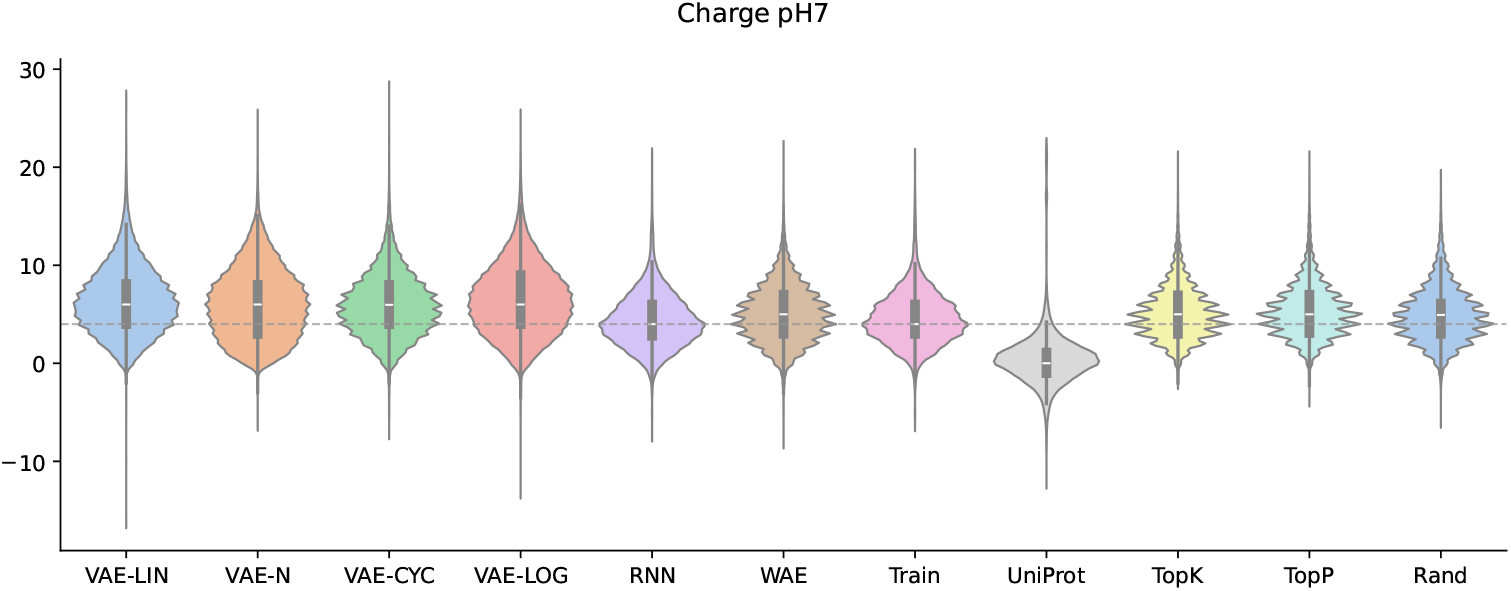
Distributions of the charge at pH 7. Distributions of the charge at pH 7 shown as violin plots including median and interquartile distances. The distributions of all randomly generated sequences, as well as training and comparison datasets are shown. The gray dashed line marks the median of the training datasets’ distribution for easier visual comparison.

### 3.2 Manifold learning

Figure 6 shows the 2D-visualization of the feature vectors, of the sequences generated by the WAE, after filtering (section 2.6), in cyan and the sequences used for training in green using UMAP as described in section 2.8. Figure 6 depicts the clustering performed by UMAP. The clustered instances show broad diversity. Instead of forming multiple or clearly separated clusters, there is one central cluster that consists of smaller subgroups or areas with a higher local point-density. Moreover, there is an absence of domains in which only a single group of instances is present. In almost all regions of the map where training data can be found, samples from the WAE can be found in the close neighborhood. The visualization of the clustering with t-SNE can be found in supplementary figure 9.

**Fig. 6.**
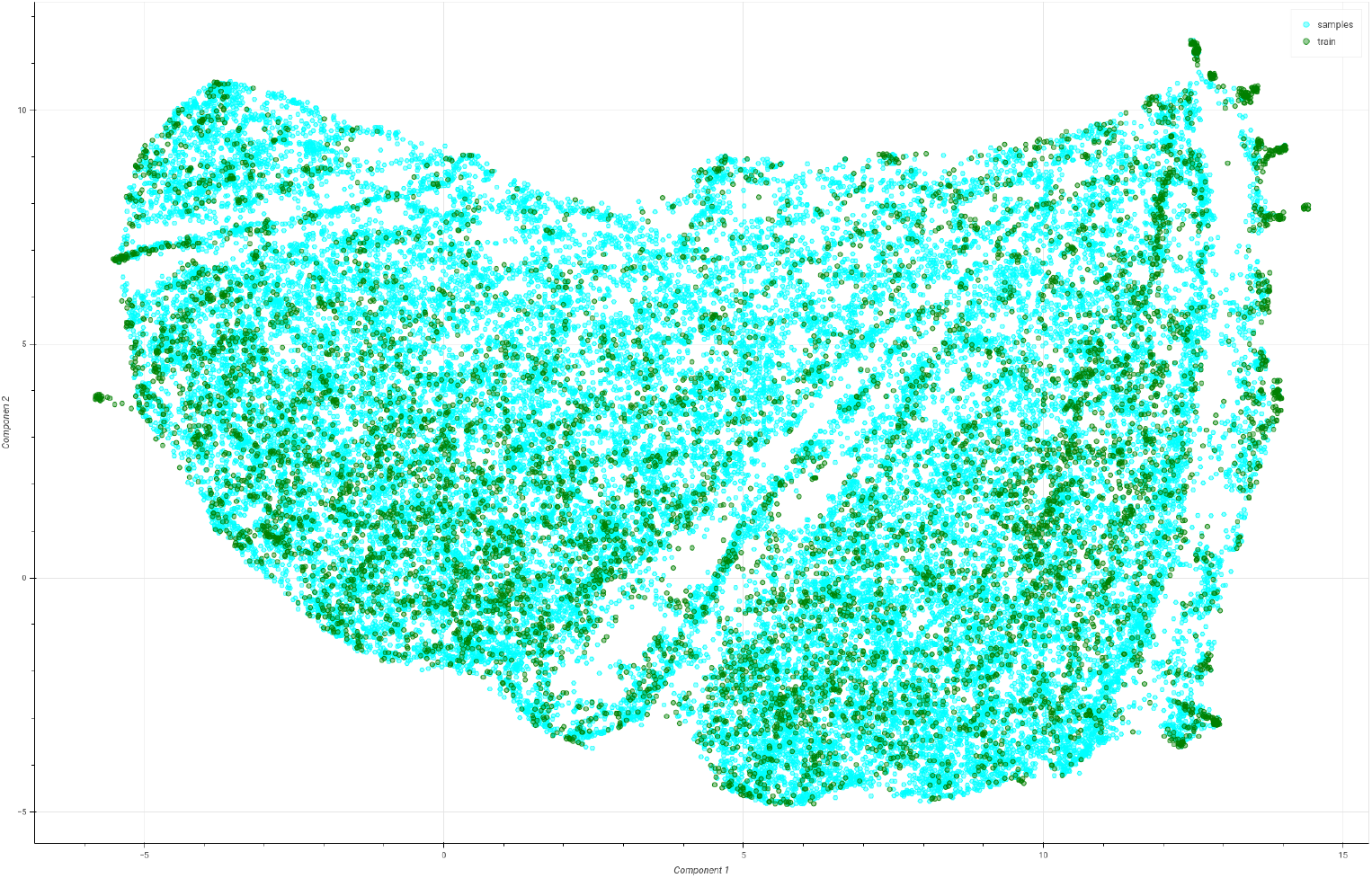
Visualization of the peptide property clustering performed by UMAP. Visualization of peptide properties calculated for the sequences generated by the WAE and the training dataset.

### 3.3 Template-based multi-sequence sampling

The results of the VAE multi-sampling process are presented in the following section. As mentioned in section 2.5, 25 samples are generated for each template sequence. Here, a comparison is made between the generated samples and their respective templates to capture sequence similarity. Subsequently, the abundance of the generated variants is analyzed. The results of these analyses are summarized in Table 1. In order to facilitate a comparison of the sequence similarity, the mean normalized Damerau-Levenshtein (LD) distance between the template sequence and its generated variants was calculated using the polars-ds library [61]. The models VAE-LIN and VAE-CYC achieve the highest distances, with 0.526 and 0.512, respectively. The mean number of unique variants generated per template ranged from 13 to 14 for all VAEs. The application of different annealing schedules has been shown to reduce the number of posterior collapsed samples, as evidenced in Table 1. Templates exhibiting posterior collapse are identified by producing only one particular sample. When an annealing schedule is applied, the number of templates for which more or equal to 20 distinct variants could be generated increases. In this case, the VAE with the linear annealing schedule performs best.

**Table 1.** Summary presenting the mean normalized Damerau-Levenshtein (LD) distance, the mean number of generated variants per template, the mean number of templates without distinct variants and the number of templates with a minimum of 20 variants.

| Auto-encoder | Mean Levenshtein distance | Mean # variants | # Collapses | $\geq 20$ |
| --- | --- | --- | --- | --- |
| VAE-CYC | 0.512 | 13 | 53 | 13 |
| VAE-LIN | 0.526 | 14 | 34 | 26 |
| VAE-LOG | 0.484 | 13 | 51 | 19 |
| VAE-N | 0.428 | 13 | 68 | 6 |

In order to compare the sequence properties of generated variants, the hydrophobicity, hydrophobic moment, and the charge at pH 7 were calculated using the peptides.py library and visualized in figure 7. For easier visualization only the templates with more or equal to 20 generated different variants from the VAE-LIN model was used in figure 7. Similar plots for the models VAE-N, VAE-LOG, and VAE-CYC are given in supplementary figures 6-8.

**Fig. 7.**
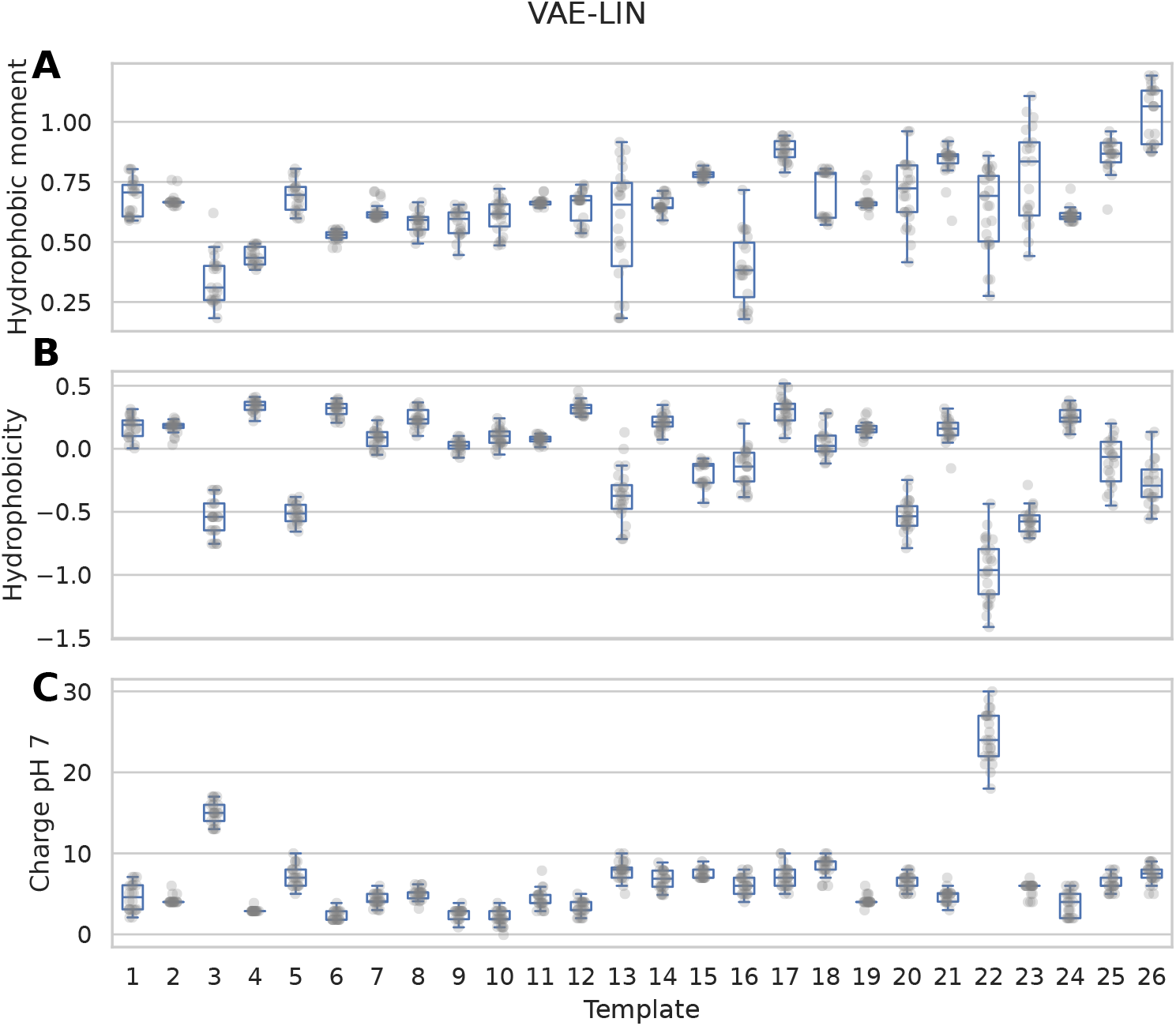
Descriptors of templates and the resulting variants of the VAE model with linear annealing. The hydrophobic moment (A), hydrophobicity (B) and charge at pH 7 (C) of the template and its generated sequences of the VAE with linear annealing. The transparent grey dots represent the single descriptor values for each generated variant of the templates 1-26, which are numbered on the x-axis.

In order to illustrate the differences between the variants generated from template sequence GSKKPVPIIYCNRRTKCQRM, an example is given in supplementary figure 10. This sequence was generated by the VAE model using the cyclic annealing schedule. In supplementary figure 10, a multiple sequence alignment done with ClustalO [62] and the biotite python library [63] between the template sequence and the generated variants is displayed. Peptide positions with varying amino acids are shown with a lighter background color.

### 3.4 Template-based sequence reconstruction

This section describes the results of the sequence reconstruction performed by the auto-encoder-based models. With each auto-encoder-based model, a single reconstruction sample was generated for all sequences from the reconstruction dataset. Table 2 presents the average reconstruction accuracy per peptide position and the number of fully reconstructed sequences. Figure 8 illustrates the reconstruction accuracy over the peptide positions per model. A Savitzky-Golay filter from the scipy library [64] was used for smoothing. The WAE demonstrates the highest overall reconstruction accuracy compared to the VAE models with a mean accuracy of 69 %. Furthermore, it produces the highest number of fully reconstructed peptides, with 111 of 720 identified. The VAE, without annealing, performs the worst of all auto-encoder-based models.

**Table 2.** Mean reconstruction accuracy per peptide position and the number of fully reconstructed sequences of each of the auto-encoder-based models.

| Model | Reconstruction accuracy | 1:1 Reconstructions |
| --- | --- | --- |
| WAE | 69 % | 111 |
| VAE-CYC | 60 % | 25 |
| VAE-LIN | 57 % | 20 |
| VAE-LOG | 56 % | 10 |
| VAE-N | 51 % | 5 |

**Fig. 8.**
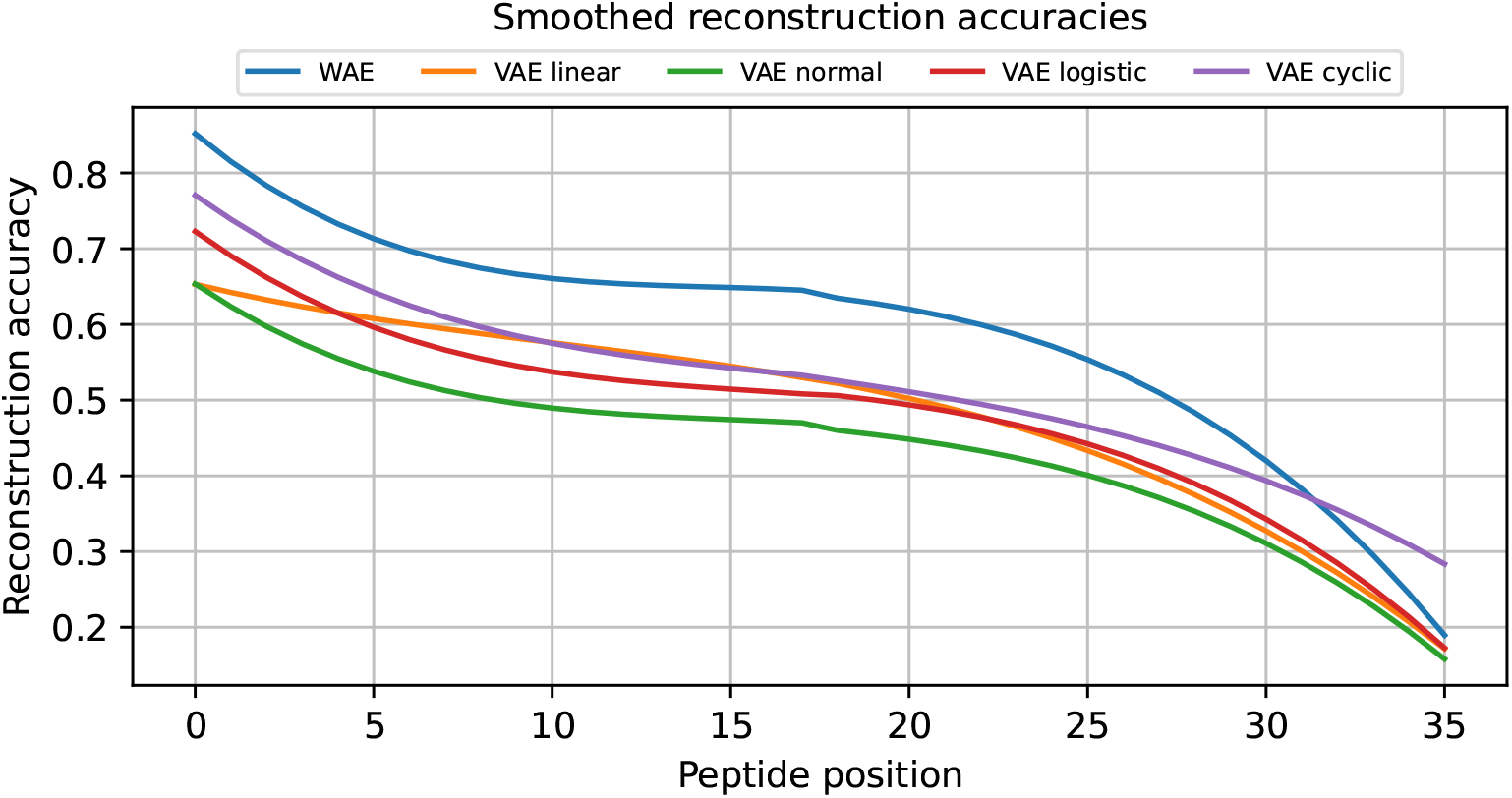
Sequence reconstruction accuracy smoothed by a Savitzky-Golay filter. The accuracy of the sequence reconstruction at each position for the five models. The accuracy of all models decreases with increasing peptide position. The WAE demonstrates high performance compared to the VAEs. Among the VAEs, the highest reconstruction accuracy was achieved with the cyclic annealing schedule.

The reconstruction accuracy achieved by the VAE models ranges from 51 % to 60 %, with a notable dependency on the annealing schedule employed. The cyclic annealing approach exhibits the highest reconstruction accuracy, with a mean of 60 %.

### 3.5 AMP prediction

This section is devoted to analyzing predictions concerning the potential activity of AMPs in the randomly generated sequences (see table 3). For this task, the AMP prediction tools: AMPScanner [54] and AntiBP3 [55], as described in section 2.7 were utilized. In addition, the two datasets with artificially generated sequences from section 2.1 were also processed, serving as a background comparison. For the initial dataset, the sequences’ length and amino acid distributions are equal to those of the training dataset. The second dataset contains only sequences generated totally at random (RT) with sequence lengths between 3 and 36 amino acids. The expectation is that the first dataset will, by chance, contain more putative AMPs than the second one. As shown in table 3, the proportion of the predicted AMPs made by AMPScanner is systematically higher than those made by AntiBP3. This disparity can be explained by the differences in the evaluation metrics values for the two prediction models reported by Bajiya *et al*. [55]. Whereas AMPScanner has a specificity of 59.0 %, a value of 90.1 % is reported for AntiBP3. Therefore, we can estimate a false positive rate of 41.0 % compared to 9.9 % for AMPScanner and AntiBP3, respectively. From this, it follows that AMPScanner has an almost fourfold higher false positive rate than AntiBP3. The mean percentage overlap of predicted sequences by the two models with a probability score ≥ 80 % is 72 %.

**Table 3.** Predictions by AntiBP3 and AMPScanner for all random generated datasets and the comparison datasets. The total number of sequences after filtering is displayed, in conjunction with the percentage of positive predictions with probability score ≥ 80 % and the overlap predicted by both tools.

| Dataset | # Sequences | $\geq 80\%$ AntiBP3 | $\geq 80\%$ AMPScanner | $\geq 80\%$ Overlap |
| --- | --- | --- | --- | --- |
| WAE | 37724 | 65.11 % | 85.08 % | 23127 |
| VAE-N | 31960 | 88.67 % | 97.83 % | 27982 |
| VAE-LOG | 28730 | 83.71 % | 96.87 % | 23730 |
| VAE-LIN | 33139 | 85.22 % | 96.23 % | 27775 |
| VAE-CYC | 35709 | 82.59 % | 94.82 % | 28737 |
| RNN | 5583 | 68.83 % | 88.75 % | 3671 |
| TopP | 32728 | 81.47 % | 90.50 % | 25063 |
| TopK | 34576 | 76.62 % | 88.73 % | 24619 |
| Rand | 34239 | 61.63 % | 79.27 % | 18788 |
| RT | 31749 | 1.06 % | 19.13 % | 191 |
| RSL | 6924 | 42.55 % | 70.83 % | 2458 |

As demonstrated in Table 3, both models accurately predict the majority of generated peptides as potential AMPs. A similar observation was reported in the study by Müller *et al*. [17] where approximately 80 % of the peptides generated with their model were classified as potential AMPs. It is important to note that Müller *et al*. reported predictions with probability score *p* ≥ 0.5 and used a different prediction model from the CAMP database [34]. As expected, the RT dataset featured the least predicted potentially active AMPs, followed by the RSL dataset, which had the second lowest number of positive predictions.

In addition to predicting active sequences in the ones generated using the non-template-based sampling methods, we searched for an exact match of the generated sequences in the AMP database DBAASP [35]. We ensured that the generated sequences were not present in the training dataset and reported only highly active AMPs annotated with a MIC value ≤ 10 µM. Table 4 presents a subset of retrieved sequences. The complete list of retrieved sequences can be found in the supplementary files. Interestingly, we find none of the sequences generated by the WAE in the databases, corroborating our hypothesis that these sequences show a higher degree of diversity than those generated with the other models and might contain sequences from undiscovered AMP families.

**Table 4.**
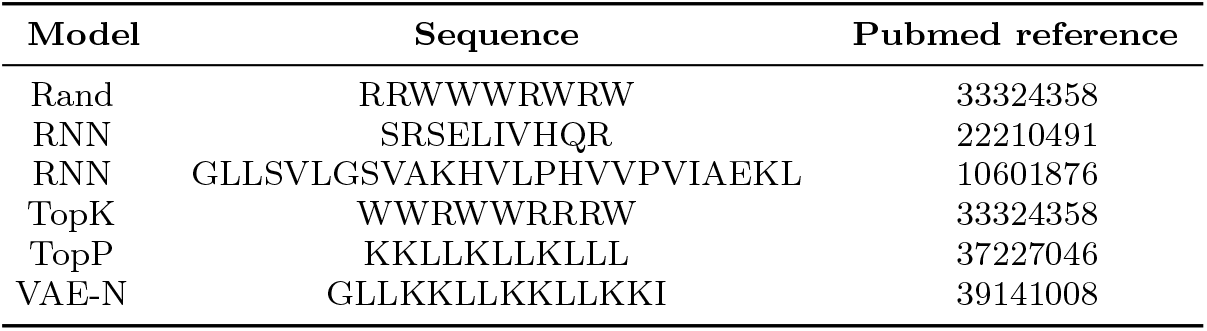
Selection of the found entries of the template free generated sequences in the DBAASP database with a reported MIC value ≤ 10 µM. The model, sequence and a PubMed reference from the DBAASP are displayed.

## 4 Discussion

In this work we compared different generative models, namely RNNs, VAEs, WAEs and a tiny version of a language model concerning their generative performance for antimicrobial peptides. Generated sequences were evaluated on sequence- and peptide property-level.

### 4.1 Sequence generation

The models for generating AMPs presented in this work, all exhibit different advantages and disadvantages. For instance, VAEs facilitate the generation of variants based on template sequences, enabling the user to generate sequences from a specific AMP family of interest (fig. 2). While the WAE can generate more diverse sequences using random sampling than VAE-based models (see section 3.1, it cannot perform template-based multi-sequence sampling (see section 2.5). Unlike the VAE-based models the RNN can only generate sequences in a random manner.

The RNN demonstrated satisfactory generative performance, even in the absence of regularization (figure 3, 4 and 5). As shown by Müller *et al*. and confirmed in this study, RNNs can learn from relatively small datasets. Müller *et al*. used around 2.000 training instances, while around 7.700 were used in this study. Therefore RNNs can be utilized in areas where experimentally validated peptide activity data is scarce. However, this model exhibits drawbacks in terms of processing speed and tends to overfit.

The LM employed in this study demonstrates medium generative performance, a phenomenon attributed to several factors. Usually, LMs are trained on natural language and extensive datasets in combination with substantial pre-training, necessitating high-performance computing hardware and long training times [47].

In contrast, the present study utilized a comparatively small language model with a limited dataset. For this reason the capacity to capture all relevant information in the training data is rather limited. Interestingly the model and training dataset have sufficient size for achieving a generative performance comparable to the VAE and WAE-models (fig. 3, 4, 5). The sampling strategies employed by the LM place considerable emphasis on character frequencies. As shown by the amino acid composition comparison in figure 3, there is a clear over-preference for the amino acid arginine (R). Notably this is the most frequently occurring amino acid in the training dataset, limiting the capacity for generating diverse sequences. This can be considered a serious drawback for this kind of generative models. Additionally, the different samplers yielded similar results regarding the peptide properties as show in figure 4. These sampling approaches may be less effective because of their strong dependence on amino acids with a high occurrence rate in the training data.

The WAE demonstrates the highest accuracy for the reconstruction task of known AMPs (see table 2). In addition, a higher proportion of the randomly generated sequences with this model pass the filtering criteria described in section 2.2 than the sequences generated by the VAEs. Furthermore, the peptides generated by the WAE were more diverse compared to those generated by VAEs because the WAE has a lower tendency to generate peptide sequences with over- or underrepresented amino acids (see fig. 3). It is conceivable that the set of randomly generated sequences by the WAE contains new unknown AMPs because the predictions made in section 2.7 were not as high as the ones for the data generated by the VAEs. This hypothesis is further substantiated by the observation that the generated sequences by the WAE exhibited a more substantial alignment with the distributions of peptide properties compared to those generated by the VAEs in section 2.6. Additionally, the analysis of the amino acid compositions between the training and generated sequences revealed a reduced disparity in the case of the WAE (see fig. 3). The VAEs also exhibited a pronounced prevalence for incorporating hydrophobic amino acids in the generated sequences. This tendency can contribute to an augmented hydrophobic moment and the preference for helical structures [65]. These structural elements are likely to be more favorably predicted by the respective models, given their enhanced similarity to the training data of the individual models or strong AMPs in general. A similar observation was reported by Müller *et al*. [17].

It is not possible to identify a single model that performs best overall. The selection of the model that fits the researcher’s use case the most, is closely tied to the application the generated peptide is designed for. For example, to select the best model, researchers should consider the type of sampling process desired. In a setting where no lead peptide sequence or family of interest available or a target organism is not specified, template-free, i.e., random sampling is naturally the way to go.

On the other hand, in case a lead peptide, a group of peptides, or a specific target organism is available, template-based sampling is recommended. In case a specific target organism is known an, appropriate starting sequence can be chosen by selecting a peptide strongly active against this organism or having desired physicochemical properties. As shown in this paper, the amino acid composition, and the relevant physicochemical properties of the template sequence are retained across the generated sequences (see fig. 7). We have also shown that these sequences exhibit a notable degree of sequence diversity (see table 1). The generated peptides are therefore expected to have a high tendency towards similar activity as the template used. Using this method, the time required for targeted AMP development can be reduced, compared to random sampling, while increasing the probability of finding sequences with the desired properties.

A further point of consideration is the selection of models or tools for the *in silico* evaluation of the generated peptides. These models may differ in training objective, architecture, or the data utilized for training. Table 3 shows that, substantial differences in AMP predictions depending on the tools used indeed exist. Moreover, conventional bioinformatics methods for AMP identification, such as alignments or sequence motif-based approaches, also yield divergent predictions [55]. At the time of writing, no commonly accepted gold standard for AMP activity prediction has been established. Although the field has seen great progress in recent years, activity prediction or even sequence generation is a very challenging task. As has been noted by Brizuela *et al*. [66, 67], both statistical and deep learning approaches perform almost equally well on these tasks, indicating that the deep learning-based models can not fully exploit their potential due to limited training dataset sizes. Specifically, the classification of AMPs is still challenging, due to their limited length and broad spectrum of activity. It is also possible that some entries in databases are erroneously annotated.

Furthermore, for some sequences, multiple strongly differing activities are reported for the same target organism. For example entry DBAASPS 4737 from the DBAASP database with MIC values reported between 4 and 32 mg*/*ml for the organism *Clostridium perfringens* ATCC 9081. Assays used for MIC determination depend on various environmental parameters such as pH, temperature or ionic strengths. These values are not always stated in the original literature and therefore also missing in the databases. This leads to entries showing high variability in reported activity values from different sources, rendering training data partially inconsistent, thereby reducing classification or regression accuracy.

### 4.2 Manifold learning

The visualization of the generated samples or their properties can be helpful with the selection process. In the case of auto-encoder-based models, this can either be accomplished by a visualization of the latent representation by t-SNE or UMAP, as previously done by Renaud *et al*. [27] and Dean *et al*. [24, 25], or by using peptide properties. Here we use a feature vector, i.e., peptide properties instead of their latent representations by the auto-encoder. Doing so, in a scenario where template-free generation is used, steering the selection process in a more template-based direction is made possible. In this case, known AMPs of interest are used as the comparison target. This is especially useful when WAE-based models are used for peptide generation because, as outlined in section 2.3, these models do not allow for template-based multi sequence generation. Nonetheless WAE-models perform better in comparison to VAEs in terms of retaining relevant properties of the training data as shown by Tolstikhin *et al*. [20]. This is in line with our findings concerning AMP properties (figure 4 and 5).

The missing separation between the clustered instances in figure 6 and supplementary figure 9 support, the previous observations, that the generated peptides share a high similarity in terms of peptide properties to the original training data. Additionally, the generated sequences show a high degree of diversity, as they can be found in almost all regions of the two-dimensional maps where training data can be found.

### 4.3 AMP prediction

*In silico* prediction of the generated peptides activity should not be used as a sole indicator for candidate selection. Depending on the use case, a multi-level comparative approach should include peptide properties or sequence similarity to a set of reference peptides of interest. Given the observation that a lower percentage of the randomly generated sequences by the WAE are predicted to be an actual AMP, we assume that the sequences generated might contain sequence pattern characteristics for AMPs that are not recognized by the prediction models. We build our hypothesis on the following observations. The sequences generated by the WAE show the fewest differences in amino acid composition of all models compared to the training dataset (fig. 3). This observation holds for both lower and higher abundant amino acids. Furthermore, fewer sequences are removed during the filtering process, i.e., sequence deduplication and the removal of sequences in which a single amino acid is repeated seven or more times in a row, indicating a higher degree of sequence diversity compared to the other models (table 3).

Additionally, the WAE reproduced the median of all the peptide descriptor distributions with the highest accuracy of all models. This is in line with the observation by Das *et al*. [1], stating that the reconstructed sequences by the WAE model were more diverse than those generated by a plain VAE. This observation should also hold for the randomly generated sequences. Since, in contrast to VAEs, WAEs force the continuous mixture *Q*_*Z*_ = ⎰*Q*(*Z* | *X*)*dP*_*X*_ to match the prior *P*_*Z*_ as described in section 2.3, latent representations of different sequences get a chance to stay farther away from each other than in the case of the VAE [20]. This allows the WAE to generate samples of higher quality for both reconstruction and random generation in both cases.

AMP prediction with AntiBP3 was also performed for the sequences generated by the VAE models using template-based multi-sequence sampling as described in section

3.3. As discussed, in section 3.5 AntiBP3 performs better than AMPScanner regarding specificity and sensitivity. Therefore, the prediction was performed using AntiBP3 only. Filtering of the generated sequences was done in three steps. First, the sequences were deduplicated, and second, sequences less than eight amino acids in length were discarded due to the length requirements of AntiBP3. Thirdly, a threshold for the Levensthein distance between the generated sequences and the respective template sequences was set to ≤ 5. The minimum proportion of sequences predicted to be active with a score ≥ 80 % was found to be 96 %.

## 5 Summary and Conclusion

The present study demonstrates that putative new antimicrobial peptides can be generated using various deep-learning models, including variational and Wasserstein auto-encoders [20], recurrent neural networks, and language models. As expected, the effectiveness of the methods used for sequence generation is shown to be model-dependent. Each model possesses distinct strengths and weaknesses. For instance, the Wasserstein auto-encoder exhibited superior accuracy reconstructing known antimicrobial peptides (see table 2). However, it cannot generate multiple samples from a single template sequence, a capability exhibited by the variational auto-encoders employed in this study (see section 2.5). Conversely, the recurrent neural network demonstrated notable efficacy, as previously reported [17]. This architecture can operate with reduced data requirements compared to other deep learning models but exhibits comparatively lower processing speeds. The auto-encoder-based models outperforms the language model utilized in this study. This might be because language models are designed to process natural languages and rely heavily on word patterns in human language. These models may not be suitable to generate short antimicrobial peptides to the same extent as auto-encoders without pre-training on a large corpus of peptide sequences.

A critical consideration is the intended use case of the generative model in combination with an appropriate sampling method. Determining whether, template-based or template-free sequence generation is appropriate for the intended application is a critical decision that must be made with careful consideration. Template-based sequence generation may be applied in cases where an active peptide with desired properties is used to generate sequences with similar properties as the template itself. On the other hand, template-free sequence generation is suitable for cases lacking lead peptides. Another point to consider is the choice of the *in silico* methods used to predict AMP activity. A plethora of tools are available for the prediction of peptide activities [68]. The models’ sensitivity and specificity depend on the datasets and peptide activity thresholds used for training. Therefore, a model that fits the use case has to be selected. Furthermore, the models utilized in this study demonstrated an ability to preserve essential peptide descriptor values (see fig. 4 and fig. 5) and amino acid sequence positions (see fig. 3) in the generated sequences. This finding serves to strengthen the hypothesis that our models possess the capacity to generate putative sequences that exhibit characteristics similar to those of known AMPs (see fig. 4, 5, 6 and supplementary figure 9). Using Manifold learning, we could support our previous findings that WAE-based generative models can generate sequences with a high degree of diversity and yet maintain relevant AMP properties.

Notably, all models employed in this study generated AMP sequences that were not utilized for training purposes. Some of these sequences are present in the DBAASP database. Among those we found AMPs with experimentally determined MIC values reported as less than ≤ 10 µM (table 4), proving that our models are indeed capable to generate highly active AMP sequences.

## Supporting information

Matched peptides in the DBAASP database

## List of abbreviations

AA: Amino Acid
AMP: Antimicrobial Peptide
AMR: Antimicrobial Resistance
ELBO: Evidence Lower Bound
GRU: Gated Recurrent Unit
IC: Inhibitory Concentration
LM: Language Model
LR: Learning Rate
LSTM: Long-Short-Term-Memory
MIC: Minimum Inhibitory Concentration
MMD: Maximum Mean Discrepancy
RNN: Recurrent Neural Network
VAE: Variational Auto-Encoder
WAE: Wasserstein Auto-Encoder

## Supplementary information

This article has two supplementary files. The file supplements.pdf contains additional figures and tables. The file matched dbaasp peptides.csv contains all generated peptides also present in the DBAASP database, that were not used for training.

## Declarations

### Ethics approval and consent to participate

Not applicable

### Consent for publication

Not applicable

### Availability of data and materials

All data and trained models are available on Zenodo. The code to train the models is available on GitHub.

- Zenodo: https://doi.org/10.5281/zenodo.17411954
- GitHub: https://github.com/devshibe/amp-autencoders

### Competing interests

The authors declare that they have no competing interests.

### Funding

This work was supported by a doctoral scholarship from the Department of Biology and Chemistry of the Justus Liebig University Giessen. This work was supported by the *Forschungscampus Mittelhessen* (FCMH, Research Campus of Central Hessen).

### Author contributions

- LB and FC Study design
- LB implementation
- LB and JH evaluation
- JH, FC, RM, AG reviewed
- All authors read and approved the manuscript

## Acknowledgements

Not applicable

## Notes

https://doi.org/10.5281/zenodo.17411954

https://github.com/devshibe/amp-autencoders

### Competing Interest Statement

The authors have declared no competing interest.

## References

[1] Das, P. et al. Accelerated antimicrobial discovery via deep generative models and molecular dynamics simulations. Nature Biomedical Engineering 5, 613–623 (2021).

[2] Naghavi, M. et al. Global burden of bacterial antimicrobial resistance 1990–2021: a systematic analysis with forecasts to 2050. The Lancet 404, 1199–1226 (2024). URL https://linkinghub.elsevier.com/retrieve/pii/S0140673624018671.

[3] Naddaf, M. 40 million deaths by 2050: toll of drug-resistant infections to rise by 70. Nature 633, 747–748 (2024).

[4] Rice, L. Federal Funding for the Study of Antimicrobial Resistance in Nosocomial Pathogens: No ESKAPE. The Journal of Infectious Diseases 197, 1079–1081 (2008). URL https://academic.oup.com/jid/article-lookup/doi/10.1086/533452.

[5] Kurbatfinski, N., Kramer, C. N., Goodman, S. D. & Bakaletz, L. O. ESKAPEE pathogens newly released from biofilm residence by a targeted monoclonal are sensitized to killing by traditional antibiotics. Frontiers in Microbiology 14, 1202215 (2023).

[6] Roque-Borda, C. A., Primo, L. M. D. G., Franzyk, H., Hansen, P. R. & Pavan, F. R. Recent advances in the development of antimicrobial peptides against ESKAPE pathogens. Heliyon 10, e31958 (2024). URL https://linkinghub.elsevier.com/retrieve/pii/S2405844024079891.

[7] Magana, M. et al. Options and Limitations in Clinical Investigation of Bacterial Biofilms. Clinical Microbiology Reviews 31, e00084–16 (2018).

[8] Olivares, J. et al. The intrinsic resistome of bacterial pathogens. Frontiers in Microbiology 4, 103 (2013).

[9] Magana, M. et al. The value of antimicrobial peptides in the age of resistance. The Lancet. Infectious Diseases 20, e216–e230 (2020).

[10] Browne, K. et al. A New Era of Antibiotics: The Clinical Potential of Antimicrobial Peptides. International Journal of Molecular Sciences 21, 7047 (2020).

[11] Fjell, C. D., Hiss, J. A., Hancock, R. E. W. & Schneider, G. Designing antimicrobial peptides: form follows function. Nature Reviews Drug Discovery 11, 37–51 (2012). URL https://www.nature.com/articles/nrd3591.

[12] Bucataru, C. & Ciobanasu, C. Antimicrobial peptides: Opportunities and challenges in overcoming resistance. Microbiological Research 286, 127822 (2024). URL https://www.sciencedirect.com/science/article/pii/S0944501324002234.

[13] Brogden, K. A. Antimicrobial peptides: pore formers or metabolic inhibitors in bacteria? Nature Reviews Microbiology 3, 238–250 (2005). URL https://www.nature.com/articles/nrmicro1098.

[14] Dijksteel, G. S., Ulrich, M. M. W., Middelkoop, E. & Boekema, B. K. H. L. Review: Lessons Learned From Clinical Trials Using Antimicrobial Peptides (AMPs). Frontiers in Microbiology 12, 616979 (2021). URL https://www.ncbi.nlm.nih.gov/pmc/articles/PMC7937881/.

[15] Sani, M.-A. & Separovic, F. How Membrane-Active Peptides Get into Lipid Membranes. Accounts of Chemical Research 49, 1130–1138 (2016-06-21). URL 10.1021/acs.accounts.6b00074.

[16] Ho, Y.-H., Shah, P., Chen, Y.-W. & Chen, C.-S. Systematic Analysis of Intracellular-targeting Antimicrobial Peptides, Bactenecin 7, Hybrid of Pleurocidin and Dermaseptin, Proline–Arginine-rich Peptide, and Lactoferricin B, by Using Escherichia coli Proteome Microarrays. Molecular & Cellular Proteomics : MCP 15, 1837–1847 (2016-06). URL https://www.ncbi.nlm.nih.gov/pmc/articles/PMC5083092/.

[17] Müller, A., Hiss, J. & Schneider, G. Recurrent Neural Network Model for Constructive Peptide Design. Journal of Chemical Information and Modeling 58, 472–479 (2018).

[18] Van Oort, C. M., Ferrell, J. B., Remington, J. M., Wshah, S. & Li, J. AMP-GAN v2: Machine Learning-Guided Design of Antimicrobial Peptides. Journal of Chemical Information and Modeling 61, 2198–2207 (2021).

[19] Tucs, A. et al. Generating ampicillin-level antimicrobial peptides with activity-aware generative adversarial networks. ACS Omega 5, 22847–22851 (2020). URL https://pubs.acs.org/doi/10.1021/acsomega.0c02088.

[20] Tolstikhin, I., Bousquet, O., Gelly, S. & Schoelkopf, B. Wasserstein Auto-Encoders (2019). URL http://arxiv.org/abs/1711.01558. ArXiv:1711.01558 [cs, stat].

[21] Chen, Q. et al. GM-pep: A high efficiency strategy to de novo design functional peptide sequences. Journal of Chemical Information and Modeling 62, 2617–2629 (2022). URL https://pubs.acs.org/doi/10.1021/acs.jcim.2c00089.

[22] Das, P. et al. PepCVAE: Semi-supervised targeted design of antimicrobial peptide sequences (2018). URL http://arxiv.org/abs/1810.07743.1810.07743 [q-bio].

[23] Pandi, A. et al. Cell-free biosynthesis combined with deep learning accelerates de novo-development of antimicrobial peptides. Nature Communications 14, 7197 (2023). URL https://www.nature.com/articles/s41467-023-42434-9.

[24] Dean, S. N. & Walper, S. A. Variational autoencoder for generation of antimicrobial peptides. ACS Omega 5, 20746–20754 (2020). URL https://pubs.acs.org/doi/10.1021/acsomega.0c00442.

[25] Dean, S. N., Alvarez, J. A. E., Zabetakis, D., Walper, S. A. & Malanoski, A. P. PepVAE: Variational autoencoder framework for antimicrobial peptide generation and activity prediction. Frontiers in Microbiology 12, 725727 (2021). URL https://www.frontiersin.org/articles/10.3389/fmicb.2021.725727/full.

[26] Hawkins-Hooker, A. et al. Generating functional protein variants with variational autoencoders. PLOS Computational Biology 17, e1008736 (2021). URL https://dx.plos.org/10.1371/journal.pcbi.1008736.

[27] Renaud, S. & Mansbach, R. A. Latent spaces for antimicrobial peptide design. Digital Discovery 2, 441–458 (2023). URL http://xlink.rsc.org/?DOI=D2DD00091A.

[28] Szymczak, P. et al. Discovering highly potent antimicrobial peptides with deep generative model HydrAMP. Nature Communications 14, 1453 (2023). URL https://www.nature.com/articles/s41467-023-36994-z.

[29] Vaswani, A. et al. Attention Is All You Need (2023). URL http://arxiv.org/abs/1706.03762. ArXiv:1706.03762 [cs].

[30] Wan, F. & De La Fuente-Nunez, C. Mining for antimicrobial peptides in sequence space. Nature Biomedical Engineering 7, 707–708 (2023). URL https://www.nature.com/articles/s41551-023-01027-z.

[31] Santos-Júnior, C. D., Pan, S., Zhao, X.-M. & Coelho, L. P. Macrel: antimicrobial peptide screening in genomes and metagenomes. PeerJ 8, e10555 (2020). URL https://peerj.com/articles/10555.

[32] King, A. M. et al. Systematic mining of the human microbiome identifies antimicrobial peptides with diverse activity spectra. Nature Microbiology 8, 2420–2434 (2023). URL https://www.nature.com/articles/s41564-023-01524-6.

[33] Maasch, J. R., Torres, M. D., Melo, M. C. & De La Fuente-Nunez, C. Molecular de-extinction of ancient antimicrobial peptides enabled by machine learning. Cell Host & Microbe 31, 1260–1274.e6 (2023). URL https://linkinghub.elsevier.com/retrieve/pii/S1931312823002962.

[34] Gawde, U. et al. CAMPR4: a database of natural and synthetic antimicrobial peptides. Nucleic Acids Research 51, D377–D383 (2023).

[35] Pirtskhalava, M. et al. DBAASP v3: database of antimicrobial/cytotoxic activity and structure of peptides as a resource for development of new therapeutics. Nucleic Acids Research 49, D288–D297 (2021).

[36] Jhong, J.-H. et al. dbAMP 2.0: updated resource for antimicrobial peptides with an enhanced scanning method for genomic and proteomic data. Nucleic Acids Research 50, D460–D470 (2022).

[37] Das, D., Jaiswal, M., Khan, F. N., Ahamad, S. & Kumar, S. PlantPepDB: A manually curated plant peptide database. Scientific Reports 10, 2194 (2020).

[38] Wang, G., Li, X. & Wang, Z. APD3: the antimicrobial peptide database as a tool for research and education. Nucleic Acids Research 44, D1087–1093 (2016).

[39] Zhao, X., Wu, H., Lu, H., Li, G. & Huang, Q. LAMP: A Database Linking Antimicrobial Peptides. PloS One 8, e66557 (2013).

[40] Bournez, C. et al. CalcAMP: A New Machine Learning Model for the Accurate Prediction of Antimicrobial Activity of Peptides. Antibiotics (Basel, Switzerland) 12, 725 (2023).

[41] The UniProt Consortium et al. UniProt: the Universal Protein Knowledgebase in 2023. Nucleic Acids Research 51, D523–D531 (2023). URL https://academic.oup.com/nar/article/51/D1/D523/6835362.

[42] Watson, M. et al. Kerashub. https://github.com/keras-team/keras-hub (2024).

[43] Kingma, D. P. & Welling, M. Auto-Encoding Variational Bayes (2022). URL http://arxiv.org/abs/1312.6114. ArXiv:1312.6114 [cs, stat].

[44] Gretton, A., Borgwardt, K. M., Rasch, M. J., Schölkopf, B. & Smola, A. A kernel two-sample test. The Journal of Machine Learning Research 13, 723–773 (2012).

[45] Hochreiter, S. & Schmidhuber, J. Long short-term memory. Neural Computation 9, 1735–1780 (1997).

[46] Cho, K., van Merrienboer, B., Bahdanau, D. & Bengio, Y. On the Properties of Neural Machine Translation: Encoder-Decoder Approaches (2014). URL http://arxiv.org/abs/1409.1259. ArXiv:1409.1259 [cs, stat].

[47] Elnaggar, A. et al. ProtTrans: Toward Understanding the Language of Life Through Self-Supervised Learning. IEEE Transactions on Pattern Analysis and Machine Intelligence 44, 7112–7127 (2022). URL https://ieeexplore.ieee.org/document/9477085/.

[48] Brandes, N., Ofer, D., Peleg, Y., Rappoport, N. & Linial, M. ProteinBERT: a universal deep-learning model of protein sequence and function. Bioinformatics 38, 2102–2110 (2022). URL https://academic.oup.com/bioinformatics/article/38/8/2102/6502274.

[49] Ferruz, N., Schmidt, S. & Höcker, B. ProtGPT2 is a deep unsupervised language model for protein design. Nature Communications 13, 4348 (2022). URL https://www.nature.com/articles/s41467-022-32007-7.

[50] Chollet, F. et al. Keras. https://keras.io (2015).

[51] Lucas, J., Tucker, G., Grosse, R. & Norouzi, M. Don’t Blame the ELBO! A Linear VAE Perspective on Posterior Collapse (2019). URL http://arxiv.org/abs/1911.02469. ArXiv:1911.02469.

[52] Fu, H. et al. Cyclical Annealing Schedule: A Simple Approach to Mitigating KL Vanishing (2019). URL http://arxiv.org/abs/1903.10145. ArXiv:1903.10145 [cs, stat].

[53] Larralde, M. peptides.py (2022). URL https://github.com/althonos/peptides.py.

[54] Veltri, D., Kamath, U. & Shehu, A. Deep learning improves antimicrobial peptide recognition. Bioinformatics (Oxford, England) 34, 2740–2747 (2018).

[55] Bajiya, N., Choudhury, S., Dhall, A. & Raghava, G. P. S. Antibp3: A method for predicting antibacterial peptides against gram-positive/negative/variable bacteria. Antibiotics 13, 168 (2024). URL https://www.ncbi.nlm.nih.gov/pmc/articles/PMC10885866/.

[56] McInnes, L., Healy, J. & Melville, J. UMAP: Uniform Manifold Approximation and Projection for Dimension Reduction. ArXiv e-prints (2018).

[57] Maaten, L. v. d. & Hinton, G. Visualizing data using t-sne. Journal of machine learning research 9, 2579–2605 (2008).

[58] McInnes, L., Healy, J., Saul, N. & Grossberger, L. Umap: Uniform manifold approximation and projection. The Journal of Open Source Software 3, 861 (2018).

[59] Poličar, P. G., Stražar, M. & Zupan, B. opentsne: A modular python library for t-sne dimensionality reduction and embedding. Journal of Statistical Software 109, 1–30 (2024). URL https://www.jstatsoft.org/index.php/jss/article/view/v109i03.

[60] Bokeh Development Team. Bokeh: Python library for interactive visualization (2025). URL https://bokeh.org/.

[61] Qin, T. Polars for data science (2024). URL https://gtihub.com/abstractqqq/polars ds extension.

[62] Sievers, F. & Higgins, D. G. Clustal Omega. Current Protocols in Bioinformatics 48 (2014). URL https://currentprotocols.onlinelibrary.wiley.com/doi/10.1002/0471250953.bi0313s48.

[63] Kunzmann, P. & Hamacher, K. Biotite: a unifying open source computational biology framework in Python. BMC Bioinformatics 19, 346 (2018). URL 10.1186/s12859-018-2367-z.

[64] Virtanen, P. et al. SciPy 1.0: Fundamental Algorithms for Scientific Computing in Python. Nature Methods 17, 261–272 (2020).

[65] Eisenberg, D., Weiss, R. M. & Terwilliger, T. C. The hydrophobic moment detects periodicity in protein hydrophobicity. Proceedings of the National Academy of Sciences of the United States of America 81, 140–144 (1984).

[66] Brizuela, C. A., Liu, G., Stokes, J. M. & De La Fuente-Nunez, C. Ai methods for antimicrobial peptides: Progress and challenges. Microbial Biotechnology 18, e70072 (2025). URL https://sfamjournals.onlinelibrary.wiley.com/doi/10.1111/1751-7915.70072.

[67] García-Jacas, C. R., Pinacho-Castellanos, S. A., García-González, L. A. & Brizuela, C. A. Do deep learning models make a difference in the identification of antimicrobial peptides? Briefings in Bioinformatics 23, bbac094 (2022). URL https://academic.oup.com/bib/article/doi/10.1093/bib/bbac094/6563422.

[68] Bárcenas, O. et al. The dynamic landscape of peptide activity prediction. Computational and Structural Biotechnology Journal 20, 6526–6533 (2022). URL https://www.sciencedirect.com/science/article/pii/S2001037022005384.

