## Additional figures and tables for "Generative models for antimicrobial peptide design: auto-encoders and beyond"

### Affiliation:



### Keywords:


### URLs:

- Zenodo
- GitHub

### Supplemental Tables

**Table 1:** Versions of the python libraries used for the implementation of the generative models and the visualization of generated peptide sequences. For the implementation of the language model some of the libraries were used with different versions, mainly due to the keras_hub library.

| Library | Version | Used for |
|---|---|---|
| **Python** | 3.11.7 | Programming |
| **Polars** | 0.20.10 | Dataframes and statistics (VAEs) |
| **Polars** | >= 1.0.0 | Language model evaluation |
| **Keras** | 2.15.0 | Deep learning models |
| **Keras** | >= 3.0.0 | Language model implementation |
| **KerasHub** | >= 0.18.0 | Language model implementation |
| **Tensorflow** | 2.15.0 | VAE model implementation |
| **Tensorflow-gpu** | 2.15.0 | GPU support |
| **Tensorflow** | >= 2.18.0 | Language model implementation |
| **tqdm** | 4.66.2 | Progress bars |
| **matplotlib** | 3.8.2 | Visualization |
| **Seaborn** | 0.13.1 | Visualization |
| **Numpy** | 1.26.3 | Required by Keras and Tensorflow |
| **peptides** | 0.3.4 | peptide properties |
| **Biotite** | >=1.0.0 | Multiple sequence alignment |

**Table 2:** Antimicrobial peptide source databases with the year of the last access or state of the database.

| Database | Last state |
|---|---|
| **CAMP** | 2023 |
| **DBAASP** | 2023 |
| **APD3** | 2023 |
| **LAMP** | from Bournez et al. 2023 |
| **dbamp** | 2022 |
| **PlantPepDB** | 2020 |

**Table 3:** Overview of the hyper parameters: batch size, learning rate, number of epochs and latent dimension size used by the generative models.

| Model | Batch size | Epochs | LR | Latent dimension |
|---|---|---|---|---|
| **WAE** | 128 | 1.000 | 0.001 | 128 |
| **VAE** | 64 | 350 | 0.001 | 64 |
| **RNN** | 128 | 100 | 0.01 | - |
| **LM** | 64 | 100 | 0.001 | - |

**Table 4:** Mean, maximum and minimum percent differences in amino acid composition between training and randomly generated sequences. The amino acid with the highest deviance is also reported.

| Model | Mean | Max | Min | Most deviating amino acid |
|---|---|---|---|---|
| **WAE** | -8.7 | 40.5 | -54.9 | Cysteine (C) |
| **VAE-CYC** | -19.4 | 66.0 | -74.0 | Tyrosine (Y) |
| **VAE-LIN** | -29.6 | 51.0 | -91.7 | Glutamic acid (E) |
| **VAE-LOG** | -18.5 | 72.8 | -86.3 | Methionine (M) |
| **VAE-N** | -27.9 | 60.0 | -92.7 | Glutamine (Q) |
| **RNN** | 1.8 | 24.0 | -14.8 | Proline (P) |
| **Random** | -1.8 | 40.2 | -23.5 | Methionine (M) |
| **TopP** | -25.8 | 56.4 | -92.1 | Methionine (M) |
| **TopK** | -24.2 | 42.0 | -95.7 | Methionine (M) |

**Table 5:** Full list of peptide properties that were calculated for the feature vectors used for manifold learning for UMAP and t-SNE. (Clickable links to the peptides.py documentation)

| Property |
|---|
| Aliphatic index |
| Boman index |
| Charge |
| Hydrophobic moment |
| Hydrophobicity |
| Isoelectric point |
| Instability index |
| Molecular weight |
| Atchley factors |
| Fasgai vectors |
| VHSE scales |
| Physical descriptors |
| Kidera factors |

**Table 6:** Mean, maximum and minimum percent differences in amino acid composition between training and randomly generated sequences. The amino acid with the highest deviance is also reported.

| Algorithm | Parameter | Value |
|---|---|---|
| **UMAP** | n_neighbors | 50 |
| **UMAP** | random_state | 42 |
| **UMAP** | n_components | 2 |
| **t-SNE** | random_state | 42 |
| **t-SNE** | n_components | 2 |
| **t-SNE** | perplexity | 100 |
| **t-SNE** | negative_gradient_method | fft |
| **t-SNE** | initialization | pca |
| **t-SNE** | metric | cosine |

### Supplemental Figures

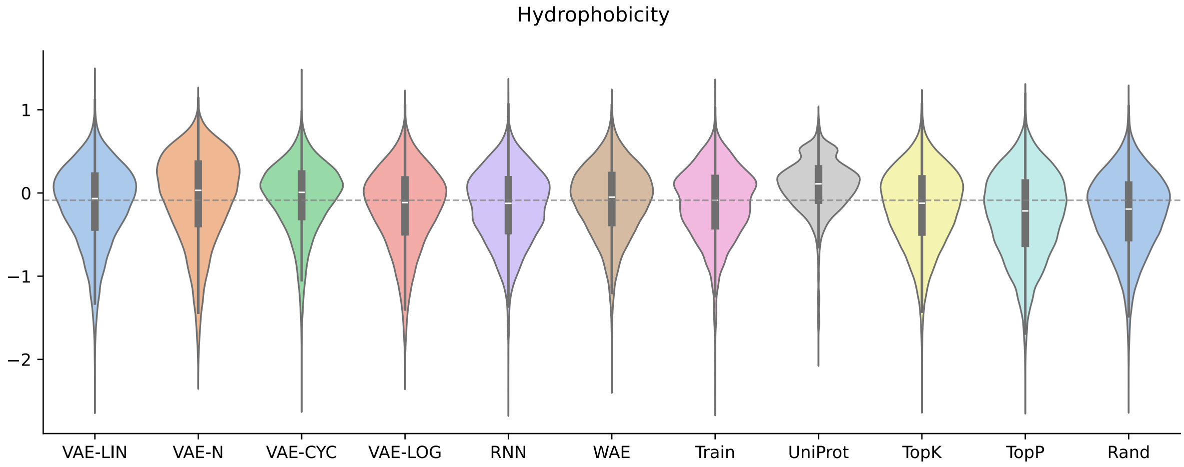


**Figure 1:** Distributions of the hydrophobicity shown as violin plots including median and interquartile distances. The distributions of all randomly generated sequences, as well as training and comparison data sets are shown. The grey dashed line marks the median of the training datasets' distribution for easier visual comparison.

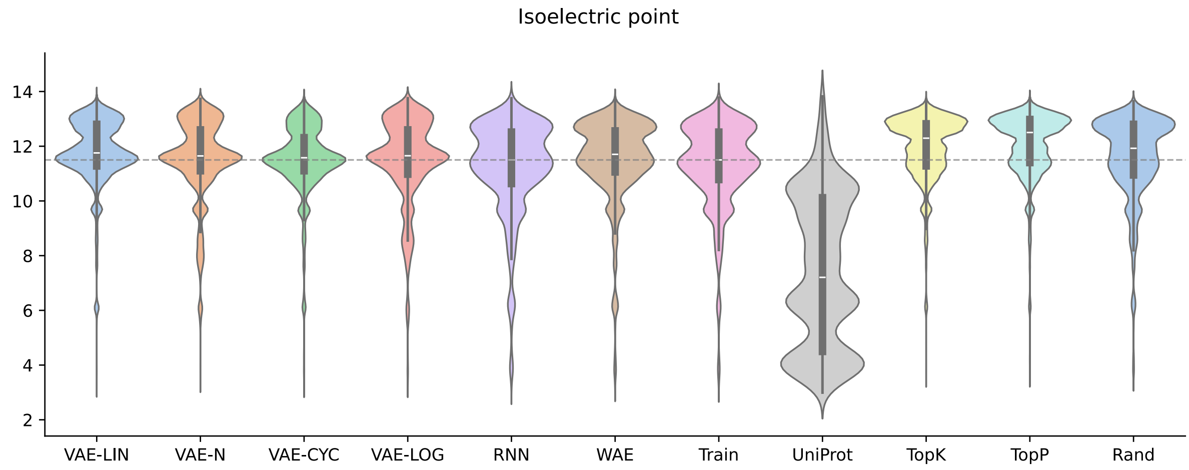


**Figure 2:** Distributions of the isoelectric point shown as violin plots including median and interquartile distances. The distributions of all randomly generated sequences, as well as training and comparison data sets are shown. The grey dashed line marks the median of the training datasets' distribution for easier visual comparison.

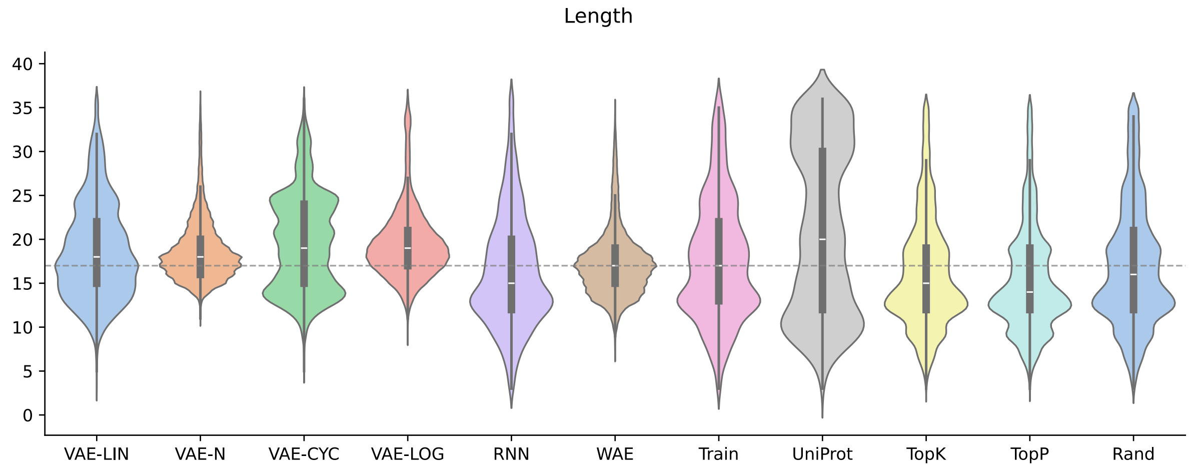


**Figure 3:** Distributions of the peptide length shown as violin plots including median and interquartile distances. The distributions of all randomly generated sequences, as well as training and comparison data sets are shown. The grey dashed line marks the median of the training datasets' distribution for easier visual comparison.

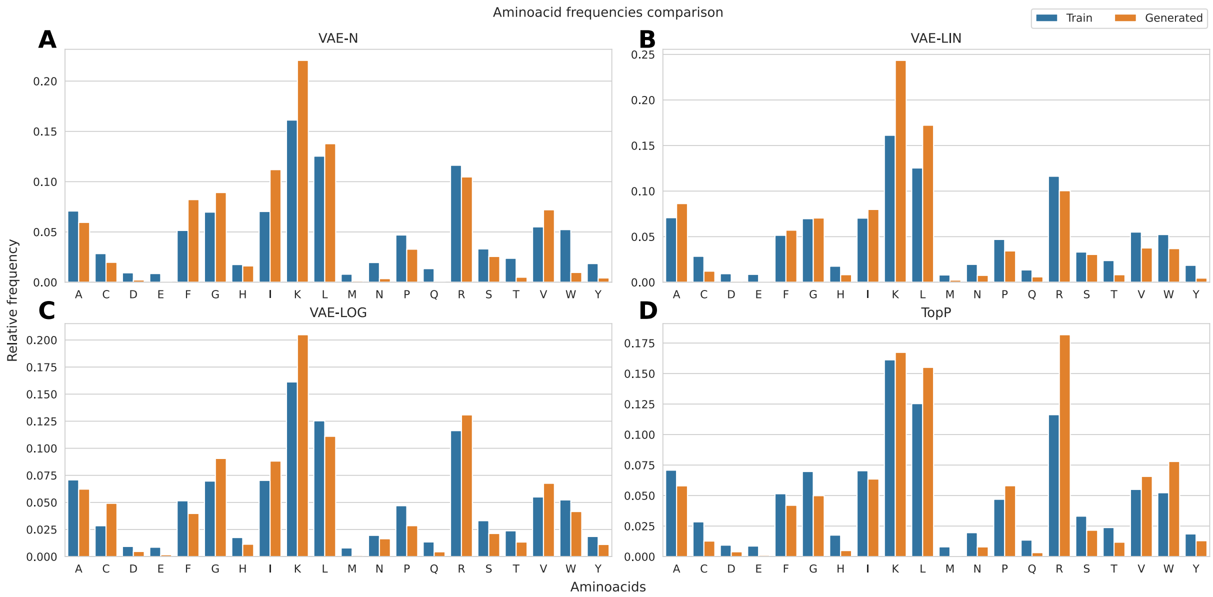


**Figure 4:** Amino acids comparison between the training set and generated sequences. Each of the sub figures shows the relative frequency of individual amino acids within the training sets (blue) and various generated sequences (orange). The results for the VAE without annealing, VAE with linear annealing, VAE with logistic annealing and TopPSampler are presented from the upper left to the lower right.

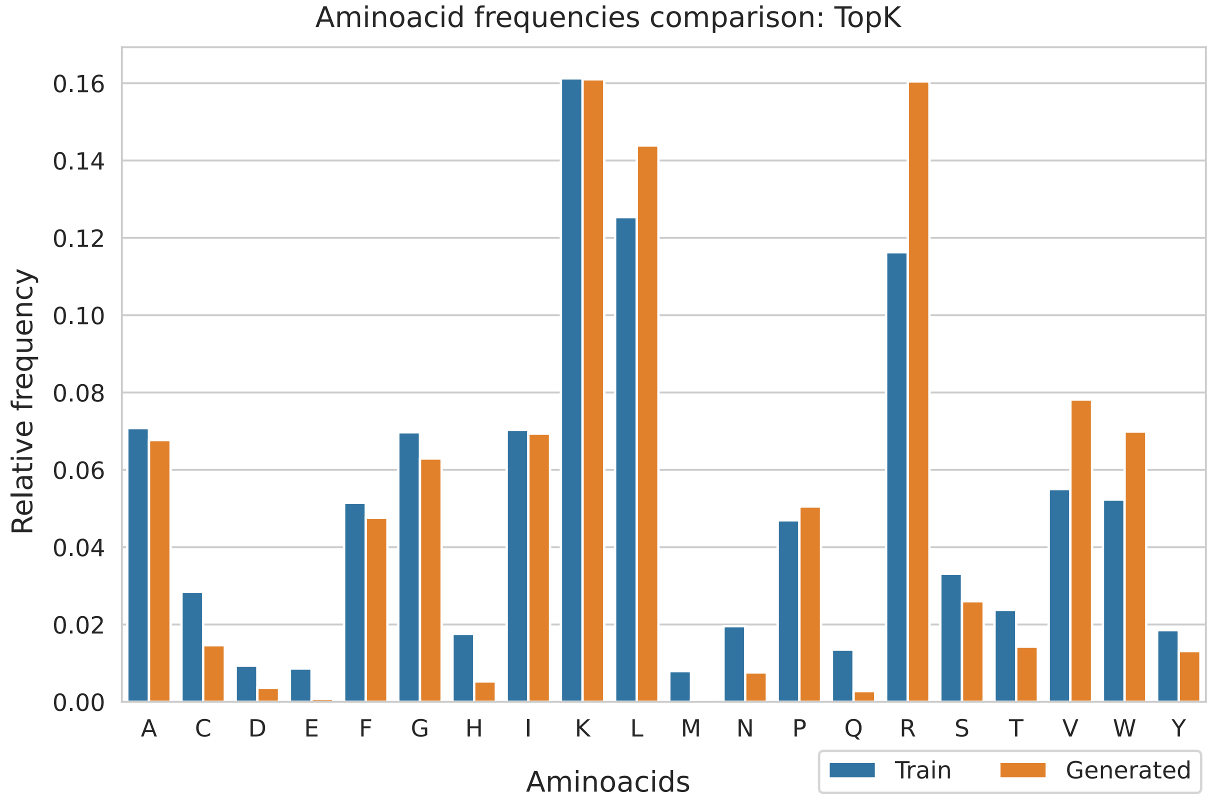


**Figure 5:** Amino acids comparison between the training set and the generated sequences of the TopK sampler. The relative frequency of individual amino acids within the training set (blue) and generated sequences (orange).

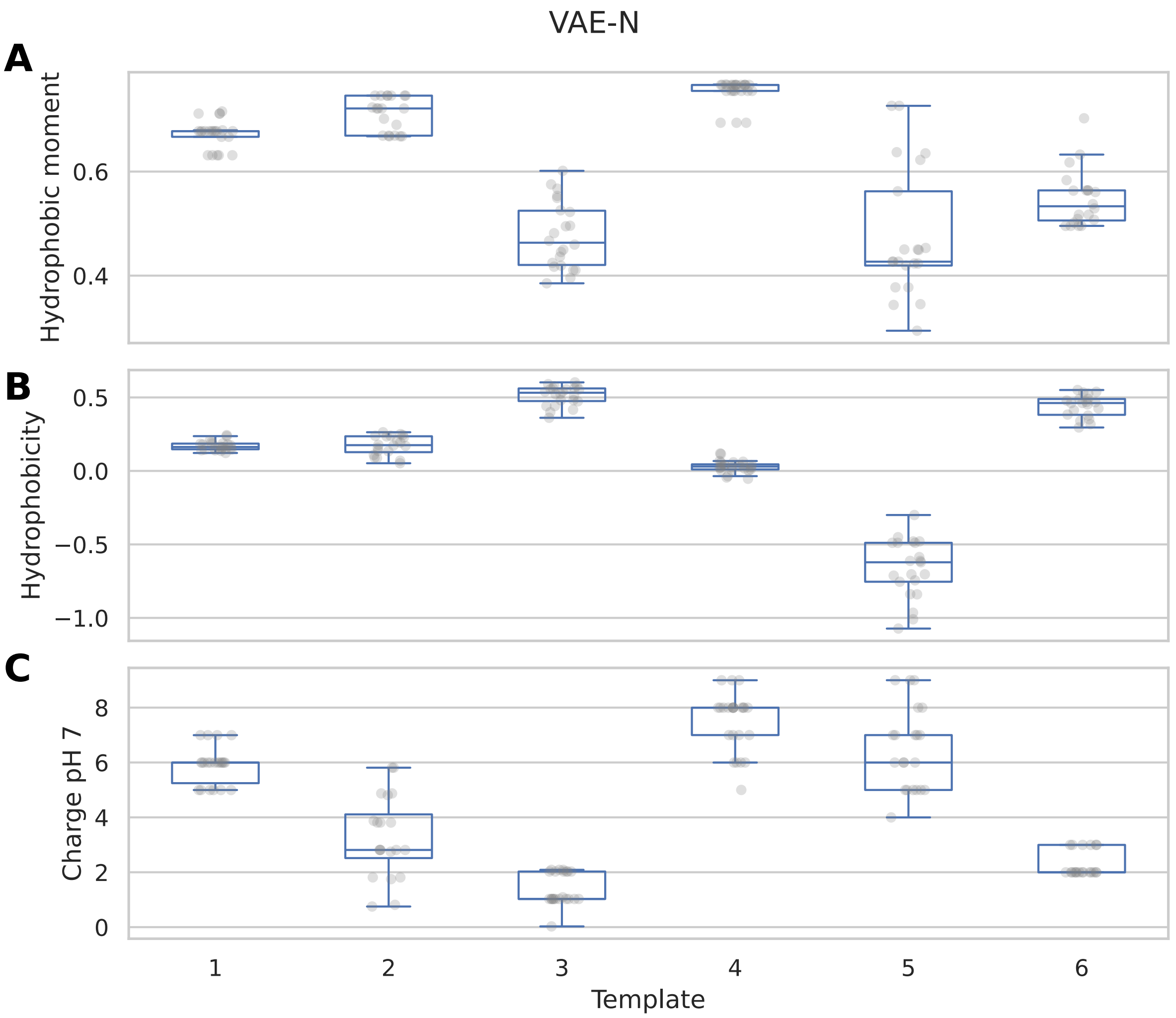


**Figure 6:** The hydrophobic moment, hydrophobicity and charge at pH 7 of the template and its generated sequences of the VAE without annealing. The transparent grey dots represent the single descriptor values for each generated variant of the templates 1-6, which are numbered on the x-axis.

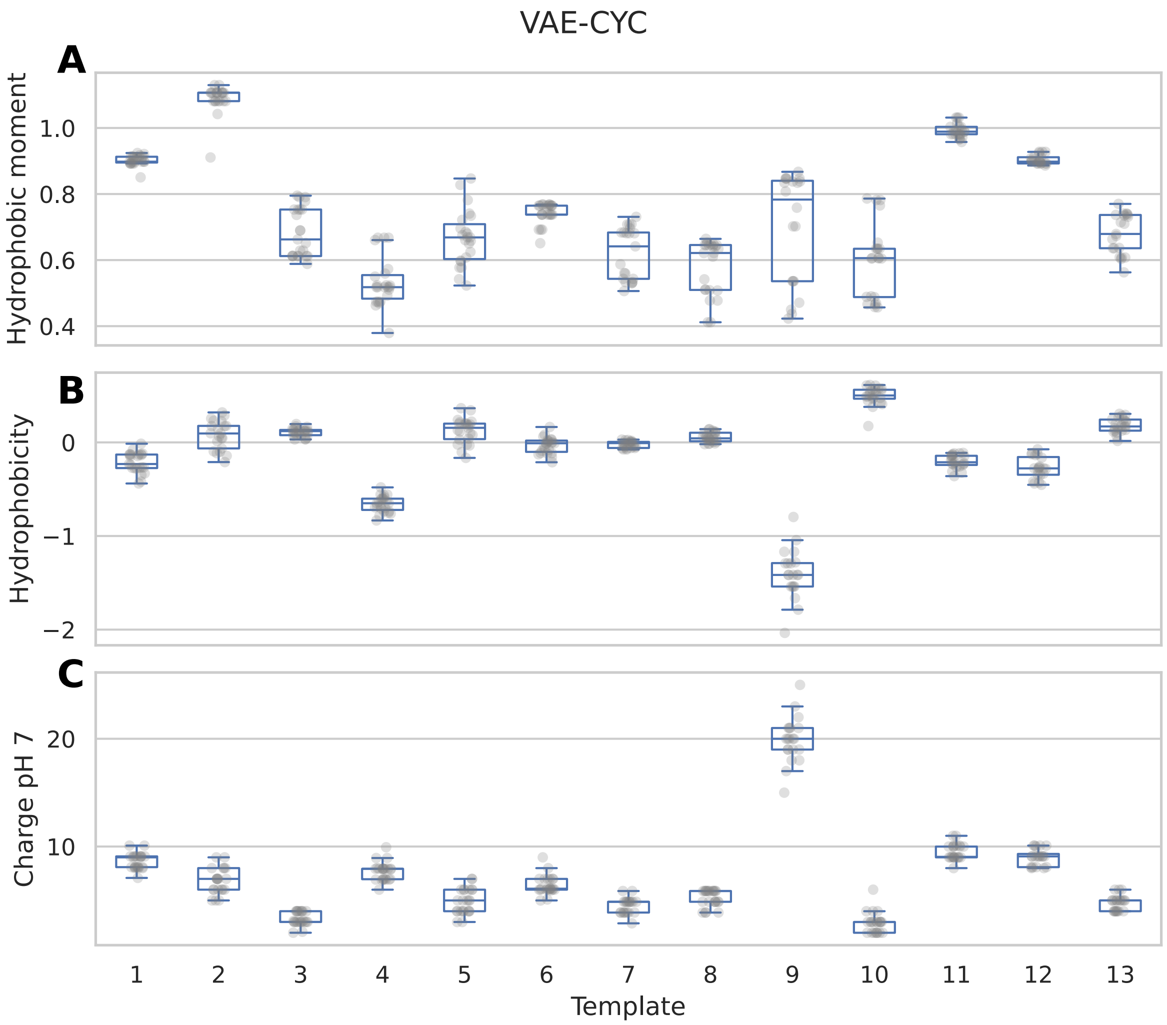


**Figure 7:** The hydrophobic moment, hydrophobicity and charge at pH 7 of the template and its generated sequences of the VAE with cyclic annealing. The transparent grey dots represent the single descriptor values for each generated variant of the templates 1-13, which are numbered on the x-axis.

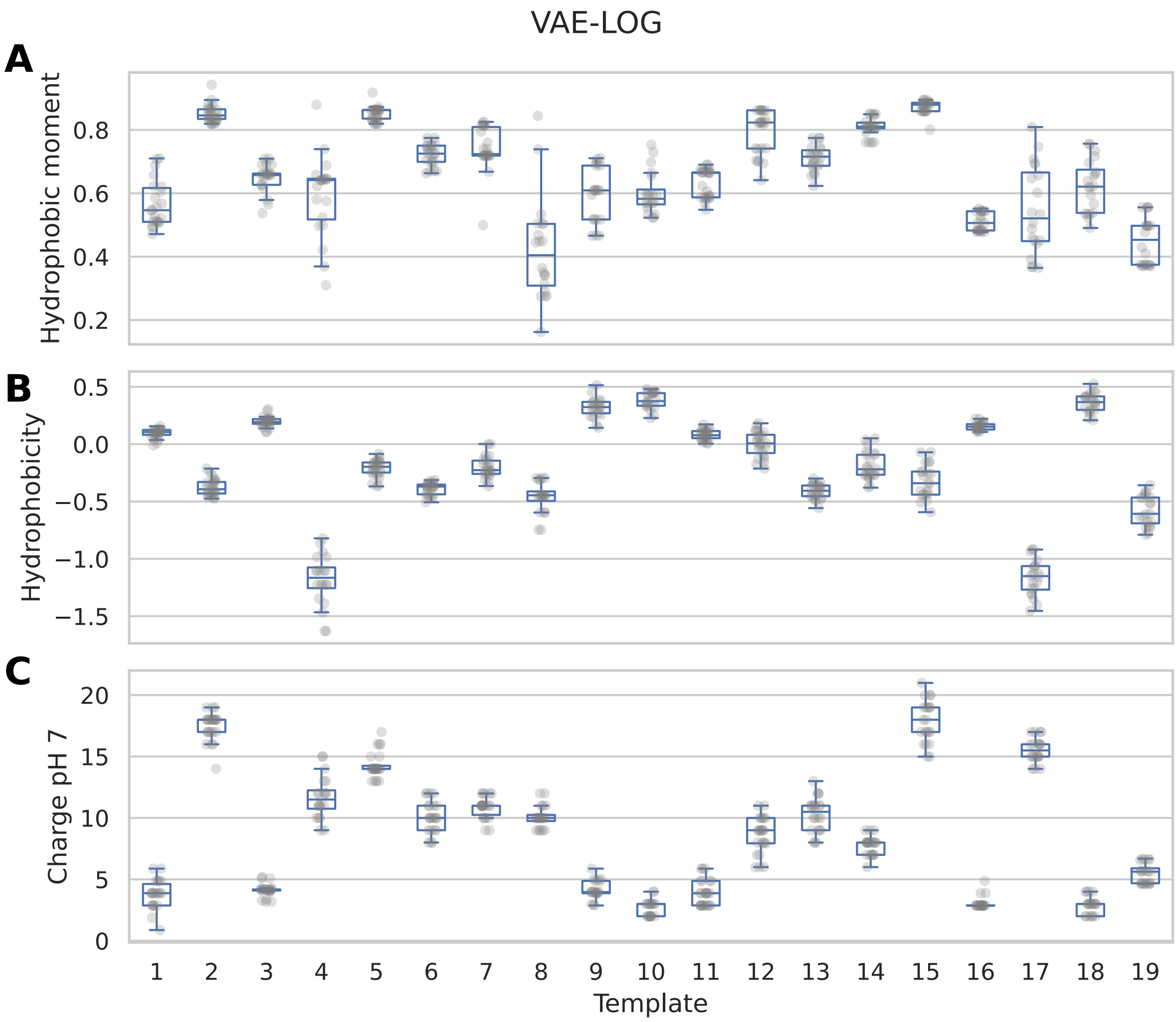


**Figure 8:** The hydrophobic moment, hydrophobicity and charge at pH 7 of the template and its generated sequences of the VAE with logistic annealing. The transparent grey dots represent the single descriptor values for each generated variant of the templates 1-29, which are numbered on the x-axis.

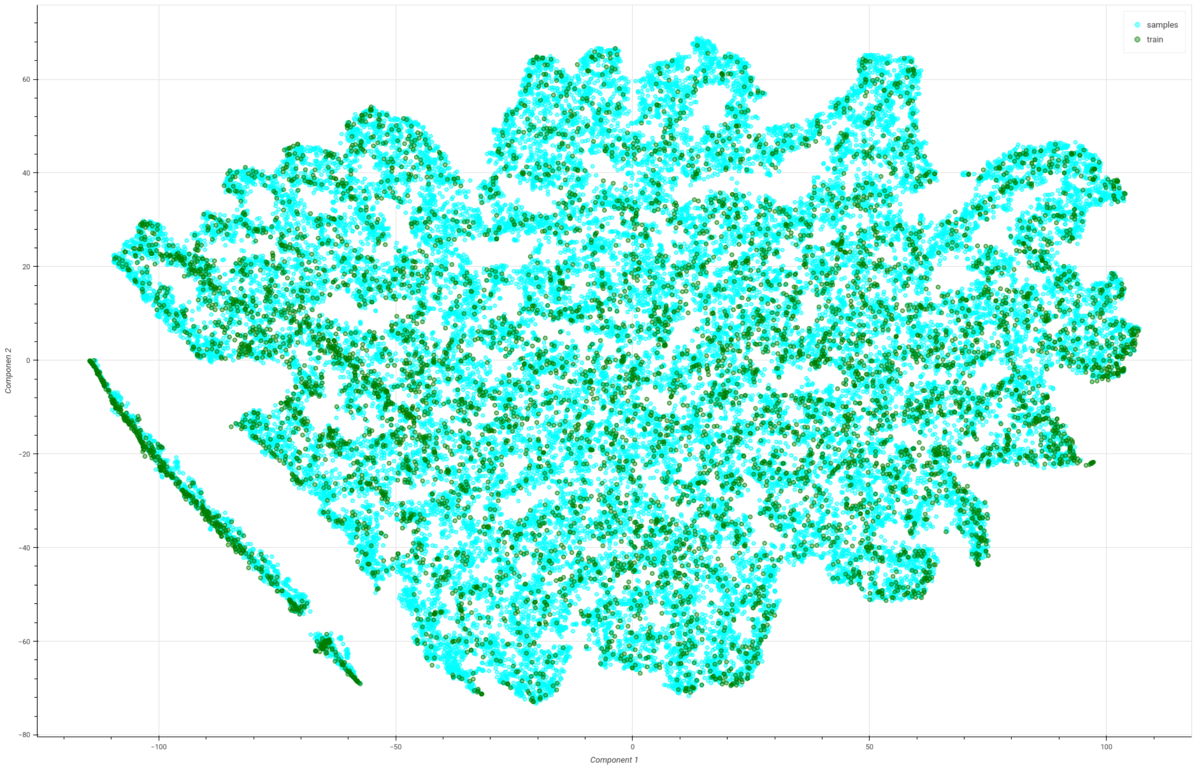


**Figure 9:** Visualization of the 2D representation of peptide feature vectors created using t-SNE.

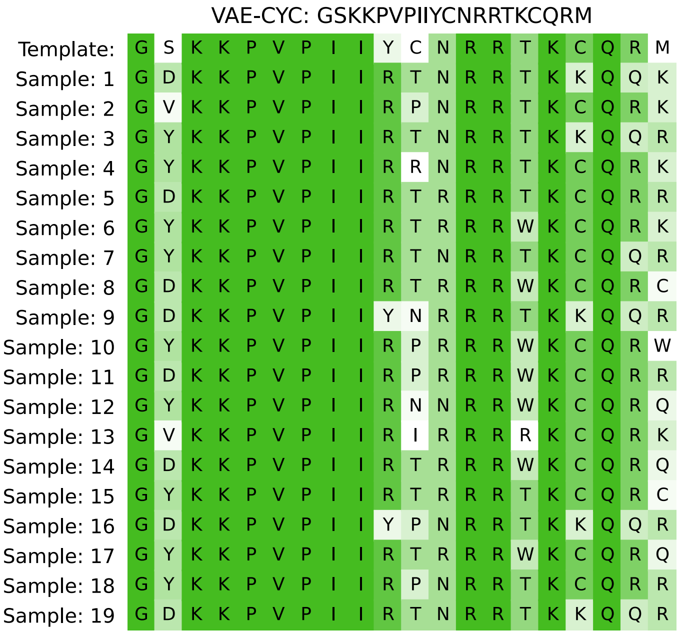


**Figure 10:** Visualization of an multiple sequence alignment from a single template sequence and the generated variants Multiple sequence alignment from the template sequence: GSKKPVPIIYCNRRTKCQRM and the generated variants by the VAE with cyclic annealing. The color corresponds to the number of variations in amino acids on the respective position in the peptide.
